# Evolutionary history of *Castanea sativa* Mill. in the Caucasus driven by Middle and Late Pleistocene paleoenvironmental changes

**DOI:** 10.1101/2023.01.11.523563

**Authors:** Berika Beridze, Katarzyna Sękiewicz, Łukasz Walas, Peter A. Thomas, Irina Danelia, Giorgi Kvartskhava, Vahid Fazaliyev, Angela A. Bruch, Monika Dering

**Affiliations:** Institute of Dendrology, Polish Academy of Sciences, Parkowa 5, 62-035, Kórnik, Poland; Adam Mickiewicz University in Poznań, Faculty of Biology, Wieniawskiego 1, Poznań, Poland; School of Life Sciences, Keele University, Staffordshire, ST5 5BG, United Kingdom; National Botanical Garden, 12 Bambis Rigi Street, Tbilisi, Georgia; Faculty of Agricultural Science and Bio-System Engineering, Georgian Technical University, Guramishvili Str. 17, Tbilisi, Georgia; Forest Development Service, Ministry of Ecology and Natural Resources of Azerbaijan, Baku, Azerbaijan; ROCEEH Research Centre, Heidelberg Academy of Sciences, Senckenberg Research Institute, Frankfurt/M, Germany; Poznań University of Life Sciences, Department of Silviculture, Wojska Polskiego 71c, 61-625, Poznań, Poland

**Keywords:** divergence time, genetic diversity, niche modelling, population structure, Early, Middle Pleistocene Transition, refugia, sweet chestnut

## Abstract

Due to global climate cooling and aridification since the Paleogene, the members of the Neogene flora were extirpated from the Northern Hemisphere or were confined to a few refugial areas. For some species, the final reduction/extinction came in the Pleistocene, but some others have survived climatic transformations up to the present. This has occurred in *Castanea sativa*, a species of high commercial value in Europe and a significant component of the Caucasian forests’ biodiversity. In contrast to the European range, neither the historical biogeography nor the population genetic structure of the species in the isolated Caucasian range has been clarified. Here, based on a survey of 21 natural populations from the Caucasus and a single one from Europe, we provide likely biogeographic reconstruction and genetic diversity details. By applying Bayesian inference, species distribution modelling, and fossil pollen data, we estimated (1) the time of the Caucasian - European divergence during the Middle Pleistocene (436.5 ka), (2) the time of divergence among Caucasian lineages, and (3) outlined the glacial refugia for species. The climate changes related to the Early Middle Pleistocene Transition and the alpine orogenic uplift in the region are proposed as the major drivers of the intraspecific divergence and European-Caucasian disjunction, while the impact of the last glacial cycle was of marginal importance.

## Introduction

Vegetation changes are key indicators of the geological and paleoclimatic evolution of the Earth. Immense geological processes throughout the Paleogene and Neogene triggered far-reaching climate and environmental transformations (Milne & Abbott, 2002; Popov *et al*., 2006). The significant legacy imprint of those changes in vegetation can be tracked down in fossil pollen records (e.g. Mahler *et al*., 2022). Moreover, a rich body of genetic evidence shows that the evolutionary histories of species, particularly the patterns of intraspecific divergence, stay closely linked to large-scale changes in their environment and thus constitute a source of peculiar knowledge about the processes of microevolution (Mayol *et al*., 2015; Fernández-López, Fernández-Cruz & Míguez-Soto, 2021; Dering *et al*., 2021).

The early Cenozoic vegetation of the Northern Hemisphere mid-latitudes consisted mainly of mixed mesophytic forests with tropical to warm-temperate taxa, which were widespread during the Paleogene and much of the Early Neogene (e.g., Kvaček & Walther, 2001; Utescher *et al*., 2007). However, during the Neogene (the Miocene – 23.3-5.33 Ma, and the Pliocene – 5.33-2.58 Ma), when pronounced cooling and aridification occurred, floristic and structural changes in the vegetation in the Northern Hemisphere were initiated. Numerous species went extinct, and a few survived in warm and humid regions of the southern latitudes and became relics currently found in a few refugia located in East Asia, west and southeast North America, and southwest Eurasia (Milne & Abbott, 2002).

The Caucasus region hosts such relict taxa in climate refugia (Tarkhnishvili, 2014), contributing to the high levels of species diversity and endemism (Myers *et al*., 2000; Nakhutsrishvili *et al*., 2017). As a result, the Caucasus is one of the global hotspots of biodiversity (Mittermeier *et al*., 2011), and yet is endangered by significant habitat loss (Mittermeier *et al*., 2011, Zazanashvili *et al*., 2004), recently further enhanced by anthropogenic climate change (Dagtekin *et al*., 2020; Dering *et al*., 2021). The Caucasus is situated at the cultural and geographical crossroads between the Black and the Caspian seas (Fig. 1). The region includes the North Caucasus (Russia), the South Caucasus (Georgia, Azerbaijan and Armenia), and extends westwards to northeastern Turkey and eastwards to northwest Iran. Therefore, the biogeographical classification of this ‘Caucasus ecoregion’ (Olson *et al*., 2001), is somewhat elusive as the geographic region overlaps with three floristic regions, i.e., Euro-Siberian, Mediterranean and Irano-Turanian, contributing to the region’s unique floristic richness and composition (Zohary, 1973; Manafzadeh, Staedler, & Conti, 2017). The Greater Caucasus belongs to the Alpine-Himalayan orogenic belt, while the Lesser Caucasus is dominated geologically by volcanic activity. The complex geology of the region, resulting from the continuous convergence of the Eurasian and African-Arabian plates (Adamia *et al*., 2011), has led to the high heterogeneity of landscape and climate, further encouraging the development, accumulation and long-term retention of biodiversity (Tarkhnishvili, 2014). The forests of Euxino-Hyrcanian provenance (Zohary, 1973), especially the Colchic vegetation at the eastern shores of the Black Sea and the Hyrcanian forests at the southern coast of the Caspian Sea, are the most emblematic vegetation units of the Caucasus ecoregion due to the high concentration of Neogene relicts, such as *Parrotia persica* (DC.) C.A.Mey. *Pterocarya fraxinifolia* (Poir.) Spach, *Zelkova carpinifolia* (Pall.) K. Koch and *Ilex colchica* Pojark. (Nakhutsrishvili *et al*., 2015). The Caucasus also is a significant evolutionary and distributional bridge for many European plant species of Asiatic origin (Kadereit, Licht & Uhnik, 2010; Jia *et al*., 2012; Volkova *et al*., 2020) and a centre of species radiation (Pokryszko *et al*., 2011; Neiber & Hausdorf, 2015; Volkova *et al*., 2020).

**Fig. 1.**
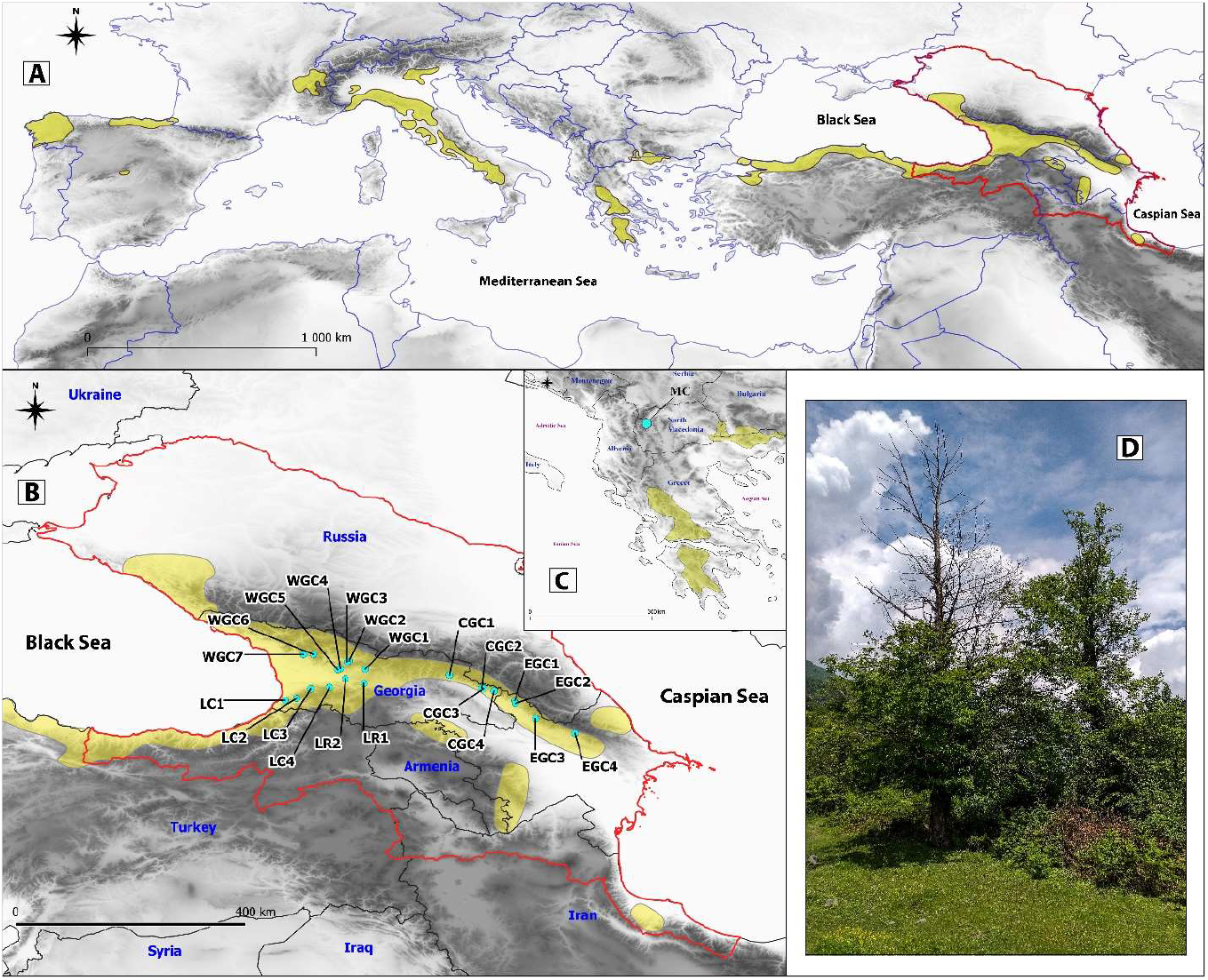
Distributional range of *Castanea sativa* (in *yellow*) according to Caudullo *et al*., (2017), (A) and throughout its entire natural range (B), in the Caucasus. Location of examined populations (B) and (C); populations acronyms as in Table 1. Ecoregional boundary of the Caucasus is delineated with a *red* line. (D) Dieback in sweet chestnut trees in Georgia possibly affected by *Cryphonectria parasitica*. Photo Credit: KS

The earliest European-Caucasian genetic splits in animal and plant populations detected using molecular data took place during the Miocene (Boroń *et al*., 2020; Levin *et al*., 2019; Tarkhnishvili, 2014) and were probably initiated by the Miocene-Pliocene global cooling and aridification (Mahler, Shatilova & Bruch, 2022; Bruch *et al*., 2011; Ivanov *et al*., 2011). Furthermore, the uplift of the Alpine-Himalayan orogenic belt since the Late Miocene accelerated, creating geographic and climatic barriers, including the Caucasus mountains (Tarkhnishvili, 2014). The range of species thus contracted and fragmented, fuelling diversification and speciation (Tarkhnishvili, 2014) and contributing to the high endemism of the region. Increased climatic variability during the Pleistocene undoubtedly reinforced these patterns (Lisiecki & Raymo, 2005). Indeed, pollen records in west Georgia (Mahler, Shatilova & Bruch, 2022) indicate that the Colchis and Hyrcania became major refugia for the Caucasian biota, especially during the last glacial period, hosting some Neogene relicts (Arpe *et al*., 2011; Shatilova *et al*., 2011; Connor & Kvavadze, 2009; Tarkhnishvili *et al*., 2012; Song *et al*., 2021).

The impact of geological and climatic changes on the genetic structure and divergence patterns remains largely unexplored for tree species growing in the Caucasus. The primary genetic patterns described involve the west-east divergence between Colchis and Hyrcanian populations (Christie *et al*., 2014; Maharramova *et al*., 2015; Sękiewicz *et al*., 2022). This genetic pattern reflects the impact of the two major refugia located in the Colchic lowland and Hyrcanian mountains during the last glaciation (Connor & Kvavadze, 2009; Tarkhnishvili, 2014; Sękiewicz *et al*., 2022). The west-east genetic split reported in *Quercus petraea* subsp. *iberica* ((M.Bieb.) Krassiln.) is attributed to the effect of the geographic barrier formed by the Likhi Range, a mountain chain running longitudinally in the Central Caucasus (Ekhavaia, *et al*., 2018). Also, this mountain range divides the region into two divergent climatic provenances – more humid to the west and dry to the east. Recently, Sękiewicz et al. (2022) reported that the geographic pattern of diversity in *Fagus orientalis* (Lipsky) was driven by this precipitation gradient. The orogeny process that resulted in the uplift of the Greater and Lesser Caucasus was a factor triggering phylogeographic patterns observed for many animal species, especially those characterized by low mobility (Tarkhnishvili, 2014). For anemophilous tree species, the results are less noticeable and rather point to pervasive gene flow in the region (Dering *et al*., 2021; Sękiewicz *et al*., 2022). Yet, the West Greater Caucasus appears as the local centre of unique genetic diversity (Ekhavaia, *et al*., 2018; Dering *et al*., 2021).

Sweet chestnut is a thermophilous tree species of high ecological, economic, and cultural importance (Conedera *et al*., 2016). It is the only *Castanea* species native to Europe and one of a small number of Neogene relict tree species present in Europe (Villani, Pigliucci & Cherubini, 1994; Dane *et al*., 2003). The species’ natural range extends from the Iberian Peninsula toward the Caspian coast in Iran (Caudullo *et al*., 2017; Conedera *et al*., 2016; Janfaza *et al*., 2017). Fossil data indicate a Paleocene-Eocene origin of *Castanea* that diversified in Central or Eastern Asia (Zhou, 1999, Lang *et al*., 2006). The genus migrated westwards through Europe to North America, the latter accompanied by further diversification (Lang *et al*., 2006). During the late Paleogene and Neogene, *Castanea* occupied much of the Northern Hemisphere (Dane *et al*., 2003), demonstrated, for example, by Eocene pollen records from Greenland (Grímsson *et al*., 2015) and the Iberian Peninsula (Barrón *et al*., 2010), as well as Miocene pollen data from Siberia (Zyryanov, 1992) and Southern California (Ballog & Malloy, 1981). In the Caucasus, fossil pollen of the genus *Castanea* is widespread in Georgia in deposits of the Late Miocene, growing at high altitudes while the lowlands were still covered by more thermophilous and hygrophilous plants (Shatilova *et al*., 2021). From Pliocene and Pleistocene deposits, *C. sativa* is commonly recorded as macrofossils and pollen from the western part of Georgia, where it grows up today (Shatilova *et al*., 2011).

*Castanea sativa* has been extensively studied within its European range (e.g. Krebs *et al*., 2004, 2019; Mattioni *et al*., 2013, 2017; Roces-Díaz *et al*., 2018; Alcaide *et al*., 2019; Pollegioni *et al*., 2020, Castellana *et al*., 2021 Fernández-López, Fernández-Cruz & Míguez-Soto *et al*., 2021), but less so in the Caucasus (Mattioni *et al*., 2017) where it might have taken refuge during the Last Glacial Maximum (LGM) (Krebs *et al*., 2004, 2019). Our understanding of the genetic ties between European and the Caucasian populations is poor. Meanwhile, the Caucasus, along with the neighbouring Asia Minor Peninsula, designate the probable initial centre of sweet chestnut domestication and hypothetical source for further dispersal to Europe that has proceeded with a cultural and economic exchange between the Middle East and Europe (Zohary *et al*., 2012; Pollegioni *et al*., 2020). Moreover, preliminary data suggest morphological distinctiveness between Hyrcanian and European populations of *C. sativa* (Yousefzadeh *et al*., 2014) such that the easternmost range of species shows distinct genetic make-up.

Here we investigate the evolutionary history of *C. sativa* in the most eastern domain of its natural distribution in the South Caucasus. Due to the marked geographic isolation and likely genetic discontinuity between the Caucasian and European populations, the South Caucasus is probably the place where natural populations of *C. sativa* with an extraordinary evolutionary past still exist. Moreover, this distinct yet unexplored gene pool is threatened by the virulent fungal parasite *Cryphonectria parasitica* (Murr.) Barr. causing chestnut blight (Dumbadze *et al*., 2018; Tavadze *et al*., 2013; Aghayeva & Harrington, 2008; Prospero *et al*., 2013), climate change (Conedera *et al*., 2021; Freitas *el al*., 2021), the newly introduced Asian chestnut gall wasp (*Dryocosmus kuriphilus* Yasumatsu, 1951, Conedera *et al*., 2021), as well as past and ongoing human pressure accompanied with a lack of active forest management (Zazanashvili *et al*., 2004). Thus, this work is an essential step to get a more complete picture of the natural history of this Neogene relict.

We applied the Approximate Bayesian Computation approach to determine the pattern of intraspecific divergence of *C. sativa* in the South Caucasus. To support our inferences regarding the potential demographic history of the species and to trace past changes in species distribution, we used Species Distribution Models (SDMs) and available paleobotanical records. With the aim of understanding the evolutionary history of this relic species, we (1) estimated the time of genetic divergence between the Caucasian and European gene pool, (2) explored the spatial pattern of differentiation in the South Caucasus, (3) reconstructed the divergence history of the species in the South Caucasus, (4) delineated the location of the probable LGM refugia of the species in the South Caucasus, and (5) reconstructed the species dynamics of the Holocene expansion in the region.

Considering the history of the Neogene flora in Europe and recent findings in *C sativa* (Fernández-López, Fernández-Cruz & Míguez-Soto, 2021), we expect to find the oldest divergence between European and the South Caucasus gene pools to be pre-Quaternary. Furthermore, we hypothesise that especially the last glacial phase triggered the intraspecific divergence in the South Caucasus gene pool of *C. sativa*. As for other species in the region (e.g. Christie *et al*., 2014), we predict a strong east-west differentiation of climate conditions, especially in terms of precipitation, and a vicariance processes in the isolated refugia of the Colchis and Hyrcania.

## Materials and methods

### Sampling, DNA extraction and genotyping

A total of 21 natural populations of *C. sativa* from the South Caucasus (17 in Georgia and four in Azerbaijan) were analysed (Table 1, Fig. 1). In addition, a single population from the North Macedonia was included to help infer the genetic distinctiveness between European and Caucasian gene pools. A total of 653 mature trees were sampled.

**Table 1.** Population identity, location, and genetic diversity parameters for studied populations from the South Caucasus and North Macedonia.

| Pop | Locality | Latitude | Longitude | Altitude | *N* | *A* | *Pa* | *Ar* | *Ho* | *uHe* | *Fis* | *Null* | Voucher<br>ID |
| --- | --- | --- | --- | --- | --- | --- | --- | --- | --- | --- | --- | --- | --- |
| LC1 | Lesser<br>Caucasus | 41.6843 | 41.8345 | 187 | 30 | 6.67 | 3 | 5.86 | 0.526 | 0.588 | 0.062 | 0.062 | NA |
| LC2 |  | 41.7287 | 42.0781 | 971 | 30 | 6.44 | 2 | 5.51 | 0.548 | 0.594 | 0.029 | 0.051 | NA |
| LC3 |  | 41.9293 | 42.3735 | 465 | 33 | 6.56 | 1 | 5.60 | 0.529 | 0.584 | 0.030* | 0.014 | NA |
| LC4 |  | 41.9581 | 42.7697 | 582 | 31 | 6.67 | 1 | 5.62 | 0.479 | 0.565 | 0.111* | 0.052 | NA |
| Average |  |  |  |  |  | 6.59 | 1.75 | 5.65 | 0.521 | 0.583 | 0.058 | 0.045 |  |
| LR1 | Likhi | 42.0437 | 43.4988 | 899 | 30 | 5.22 | 1 | 5.60 | 0.474 | 0.546 | 0.086* | 0.050 | NA |
| LR2 | Range | 42.1429 | 43.1068 | 365 | 30 | 5.78 | 0 | 4.42 | 0.483 | 0.587 | 0.081 | 0.057 | NA |
| Average |  |  |  |  |  | 5.50 | 0.50 | 5.01 | 0.479 | 0.567 | 0.084 | 0.054 |  |
| WGC1 | Western<br>Greater<br>Caucasus | 42.3441 | 43.5264 | 595 | 30 | 5.33 | 2 | 4.62 | 0.506 | 0.530 | 0.025 | 0.040 | KOR 53913 |
| WGC2 |  | 42.5205 | 43.1896 | 756 | 29 | 5.00 | 0 | 4.58 | 0.492 | 0.539 | 0.018 | 0.055 | NA |
| WGC3 |  | 42.5038 | 43.1462 | 775 | 33 | 5.56 | 0 | 4.80 | 0.529 | 0.546 | 0.033 | 0.017 | NA |
| WGC4 |  | 42.3557 | 42.9991 | 574 | 31 | 4.78 | 0 | 4.20 | 0.503 | 0.554 | 0.029 | 0.077 | KOR 53916 |
| WGC5 |  | 42.3329 | 42.9421 | 537 | 28 | 6.44 | 5 | 5.64 | 0.583 | 0.562 | 0.021 | 0.018 | KOR 53912 |
| WGC6 |  | 42.6569 | 42.4341 | 913 | 29 | 5.78 | 0 | 4.92 | 0.437 | 0.533 | 0.131* | 0.077 | NA |
| WGC7 |  | 42.6548 | 42.2216 | 343 | 31 | 6.22 | 0 | 5.39 | 0.482 | 0.518 | 0.073* | 0.024 | NA |
| Average |  |  |  |  |  | 5.59 | 1.0 | 4.88 | 0.505 | 0.540 | 0.047 | 0.044 |  |
| CGC1 | Central<br>Greater<br>Caucasus | 42.2226 | 45.3037 | 806 | 18 | 4.67 | 2 | 4.64 | 0.549 | 0.532 | 0.030 | 0.017 | NA |
| CGC2 |  | 41.9472 | 45.9865 | 679 | 30 | 4.44 | 0 | 3.88 | 0.456 | 0.471 | 0.029 | 0.020 | NA |
| CGC3 |  | 41.9326 | 46.0194 | 559 | 27 | 5.00 | 0 | 4.38 | 0.413 | 0.511 | 0.136* | 0.060 | KOR 53914 |
| CGC4 |  | 41.8854 | 46.2457 | 691 | 22 | 3.56 | 0 | 3.65 | 0.433 | 0.415 | 0.026 | 0.005 | NA |
| Average |  |  |  |  |  | 4.42 | 0.50 | 4.14 | 0.463 | 0.482 | 0.055 | 0.026 |  |
| EGC1 | Eastern<br>Greater<br>Caucasus | 41.6796 | 46.6864 | 963 | 33 | 4.56 | 0 | 3.85 | 0.473 | 0.499 | 0.084* | 0.027 | KOR 53905 |
| EGC2 |  | 41.6194 | 46.6990 | 670 | 31 | 4.11 | 0 | 3.66 | 0.436 | 0.510 | 0.082 | 0.044 | KOR 54819 |
| EGC3 |  | 41.2976 | 47.1249 | 852 | 29 | 3.56 | 0 | 3.20 | 0.481 | 0.491 | 0.019 | 0.046 | KOR 53883 |
| EGC4 |  | 40.9883 | 47.9544 | 1058 | 41 | 3.89 | 1 | 3.24 | 0.473 | 0.491 | 0.030 | 0.022 | KOR 53840 |
| Average |  |  |  |  |  | 4.03 | 0.25 | 3.49 | 0.466 | 0.498 | 0.054 | 0.035 |  |
| MC | Europe | 41.7247 | 20.8432 | 953 | 27 | 5.22 | 7 | 4.57 | 0.651 | 0.652 | 0.015 | 0.045 | KOR 53933 |

Pop – population identity, $N$ – number of surveyed individuals, $A$ – mean number of alleles, $Pa$ – number of private alleles, $Ar$ – allelic richness, $Ho$ – observed heterozygosity, $uHe$ – unbiased estimation of expected heterozygosity, $Fis$ – inbreeding coefficient, $*$ – inbreeding is an important factor in the population, $Null$ – frequencies of null alleles

DNA was extracted from dry leaves using the CTAB protocol (Dumolin *et al*., 1995). Initially, 22 polymorphic nuclear microsatellite markers (nSSRs) designed for European chestnut were chosen for study (Marinoni *et al*., 2003; Buck *et al*., 2003). However, due to amplification problems nine nSSR - CsCAT1, CsCAT6, CsCAT14, CsCAT15, CsCAT41 (Marinoni *et al*., 2003) and EMCs2, EMCs13, EMCs15, EMCs22 (Buck *et al*., 2003) - were employed in the final analyses. Among all selected nSSR loci, eight were polymorphic in all tested populations, while locus EMCs2 was polymorphic for the Macedonian population. PCR conditions were chosen according to Marinoni *et al*. (2003) and Buck *et al*. (2003). PCR products were run on Applied Biosystems™ ABI PRISM 3130 XL genetic analyzer (Thermo Fisher Scientific, USA) using an internal standard GeneScan 500LIZ^®^ (Thermo Fisher Scientific, USA). Then, fragment sizes were scored using GeneMapper 4.0 software (Thermo Fisher Scientific, USA). Finally, Raw2Gen software (IJ Chybicki, unpubl. data) was used for allele binning.

### Genetic diversity and differentiation

We calculated linkage disequilibrium (LD) in Genepop v 4.7.5 (Raymond & Rousset, 1995; Rousset, 2008) using the log-likelihood ratio statistic test for each pair of loci with Bonferroni correction. Next, the same software was used to calculate deviations from the Hardy-Weinberg equilibrium for each population and each loci. Null allele frequency was estimated using the Maximum Likelihood method implemented in ML - Null Freq (Kalinowski & Taper, 2006). Finally, GenAlEx 6.51b2 (Peakall & Smouse, 2012) was used to calculate genetic diversity parameters, including the mean number of alleles (*A*), the number of private alleles (Pa), observed heterozygosity (*Ho*) and unbiased estimation of expected heterozygosity (*uHe*). Allelic richness (*Ar*) was estimated in FSTAT 2.9.4 (Goudet, 2003).

Additionally, we tested the statistical difference in parameters of genetic structure (*He*, *Ar*, *Fis* and *F_ST_*) between different sets of populations: (1) among LC, LR, WGC, CGC and EGC populations (for abbreviations, refer to Table 1) to search for sub-regional patterns; (2) between Western Georgian populations located in the Greater and Lesser Caucasus (WGC/LC) excluding populations from the Likhi Range (LR1, LR2) to search for diversity pattern specific to different mountain ranges, and (3) between populations on the west (LC, LR, and WGC) and on the east (CGC and EGC) of the region to search for differences in diversity patterns among populations that likely derive from Colchis and Hyrcanian refugial areas, respectively. Statistical significance was calculated in FSTAT 2.9.4 based on 9,999 permutations.

Finally, we estimated the inbreeding coefficient (*Fis*) including “null alleles” correction based on the individual inbreeding model (IIM) using the Bayesian approach implemented in INEst 2.2 (Chybicki & Burczyk, 2009). Parameters were specified as 5 ×10^5^ MCMC iterations, storing every 200th value with burn-in steps of 5 x 10^4^ Two independent runs were done for each population selecting different models according to the Deviance Information Criterion (DIC), i.e., the inbred population model (*nfb*, full model where *n* null alleles, *f* inbreeding coefficient, *b* genotyping failure) and the random mating model (*nb*). Then, to estimate the extent of divergence among the Caucasian populations and regions, Wright’s fixation index (*F_ST_*) was calculated using FreeNA (Chapuis & Estoup, 2007) with “Excluding Null Alleles” (ENA) correction for the presence of null alleles. Finally, significance was tested using bootstrapping resampling over loci method with 10^4^ replications.

### Range-wide population structure

The Bayesian clustering approach implemented in STRUCTURE 2.3.4 (Pritchard, Stephens & Donnelly, 2000) was used to infer the genetic structure of the populations. This analysis was performed in two variants. First, we performed genetic clustering for the 21 Caucasian populations and a single one from the North Macedonia to preliminary explore the differentiation between the Caucasian and European gene pool. Sampling only a single site in Europe will deliver very coarse results in terms of the complexity of the genetic structure of the European and Caucasian gene pools, hence, the results of these analyses should be treated with caution. To gain a deeper insight into the wide-range genetic structure of *C. sativa* in the South Caucasus, STRUCTURE was run only for the Caucasian populations. For both variants of analysis, we kept the same conditions as follows: (1) admixture and correlated allele frequencies models (2) 10 independent runs for each K ranging from 1 to 23 (the first variant) or 1 to 22 (the second variant) (3) 10^5^burn-in periods followed by 1 x 10^6^ MCMC iterations. To infer the optimal number of clusters that best tune with our dataset, the Best K, we employed different approaches to infer K number, as commonly acknowledged by the research community (e.g., Meirmans, 2015; Cullingham *et al*. 2020). We used Evanno’s ΔK approach (Evanno, Regnaut & Goudet 2005) and log probability of the data (Ln Pr (*X|K*); Pritchard *et al*., 2000), and the algorithm based on the median membership coefficient (Q) recently proposed by Puechmaille (2016) which appears to be more accurate and reliable than commonly applied methods (Puechmaille, 2016). To obtain the K-selection plots, we used StructureSelector (Li & Liu, 2018), while CLUMPAK (Kopelman *et al*., 2015) was used to summarise and visualise the replicate runs. Being aware of the complexity of factors responsible for structuring the diversity, we interpreted the number of genetic clusters in terms of the delivered biogeographic information delivered, as it is advised by some authors (e.g., Meirmans, 2015, Cullingham *et al*. 2020). To visualise the spatial genetic structure, the mean membership coefficient (Q) values were interpolated across the landscape using QGIS 3.16.3 ‘Hannover’ (QGIS Development Team 2021) using the Inverse Distance Weighting (IDW) interpolation.

To more deeply explore the spatial pattern of the population differentiation, we used the geogenetic approach implemented in SpaceMix, which is a statistical method that allows for the visualisation of complex histories of gene flow among populations, including inferring the longdistance dispersal of genes (LDD; Bradburd, Ralph, & Coop, 2016). Briefly, this approach uses a Bayesian framework to reconstruct the genetic relatedness among populations which is shown on *geogenetic map* (Bradburd, Ralph, & Coop, 2016). The geographic location of the populations of such maps is modelled by the impact of the gene flow that affects pairwise genetic similarity. Therefore, population locations on the geogenetic map reflect more the genetic proximity rather than geographic one. Additionally, since the allele frequency covariance is a decreasing function of geogenetic distance, the LDD is manifested as an abnormally strong covariance over long geographic distances. This method shares the advantages of the non-model approach such as PCA but deals much better with genetic admixture that frequently takes place in natural populations. In SpaceMix, we used frequency allele data and tested a model that estimates the population locations and admixture (‘source_and_target’). This model locates the studied populations in the geo-genetic space independently of their true geographic locations and, at the same time, estimates the gene flow. Simulations were run using 10 initial fast runs, each for 10^5^ MCMC iterations and one long run with 10^6^ iterations with a sampling of every 10^3^. The inferred geogenetic location of the populations and their admixture sources were superimposed on the observed population sampling locations. The uncertainty in locations of the populations was visualised with 50% of credible ellipses and inferences about the source of admixture were made for each population to detect the pattern of admixture and the possibility of LDD. The overall performance of MCMC was validated by exploring the posterior probability trace, while the ability of the model to describe our data was evaluated by comparison of parametric vs. observed covariance matrix.

### Demographic history

For a probabilistic analysis of alternative hypotheses for the history of divergence of sweet chestnut lineages from the South Caucasus, we used the approximate Bayesian computation (ABC) procedure combined with supervised machine learning – Random Forest algorithm implemented in R application DIYABC Random Forest v1.2.1, hereafter ABC-RF (Collin *et al*., 2021). The STRUCTURE’s results (Supporting information 3, Fig. S2, S4) conducted on the Caucasian stands and the single European one served as a justification for assumptions about the genetic distinctiveness of the North Macedonian site, which was always assumed to be the earliest point of the divergence in tested demographic scenarios. The demographic history of the Caucasian populations was based on STRUCTURE’s K=4 performed for this set of populations (see Results). For dating the divergence points among genetic lineages, we selected individuals that showed at least 80% of ancestry to the respective cluster, hereafter called a lineage. As a result, the gene pool representing the Lesser Caucasus (Lineage I) consisted of 62 individuals, the gene pool of the West Greater Caucasus (Lineage II) consisted of 72 individuals, the Central Greater Caucasus (Lineage III) consisted of 65 individuals, and the population EGC4 that represents the most easterly located and most genetically distant Caucasian gene pool consisted of 40 individuals (Lineage IV). The European gene pool consisted of 27 individuals (Lineage V).

Seven scenarios regarding the topology of population ancestry were constructed based on the patterns of genetic differentiation revealed by STRUCTURE (Fig. 3). Based on the results of the phylogenetic studies (Janfaza *et al*., 2015), we made an assumption about the common ancestor for all genetic lineages investigated here. A single scenario included an admixture event, while the remaining six scenarios presented divergence events of different hierarchies. Scenario 1 assumes the first split at *t2* generations ago among Lineage I (Lesser Caucasus), Lineage IV (Eastern Greater Caucasus - population ECG4) and Lineage V (Europe). Further, at *t1* generations ago, Lineage II (Greater Caucasus) diverged from Lineage I. The last event included the admixture at *ta* generations ago between Lineage II and IV that resulted in Lineage III. Scenario 2 presents the differentiation of all genetic lineages throughout a single divergence event *t1* generations ago. Scenario 3 refers directly to the K=4 structure and west-east differentiation and shows that Lineages 2 and 4 diverged from Lineages 1 and 3, respectively. Finally, scenarios 4-7 assume several topologies which describe different hierarchies of divergence and ancestry for the current lineages.

**Fig. 2.**
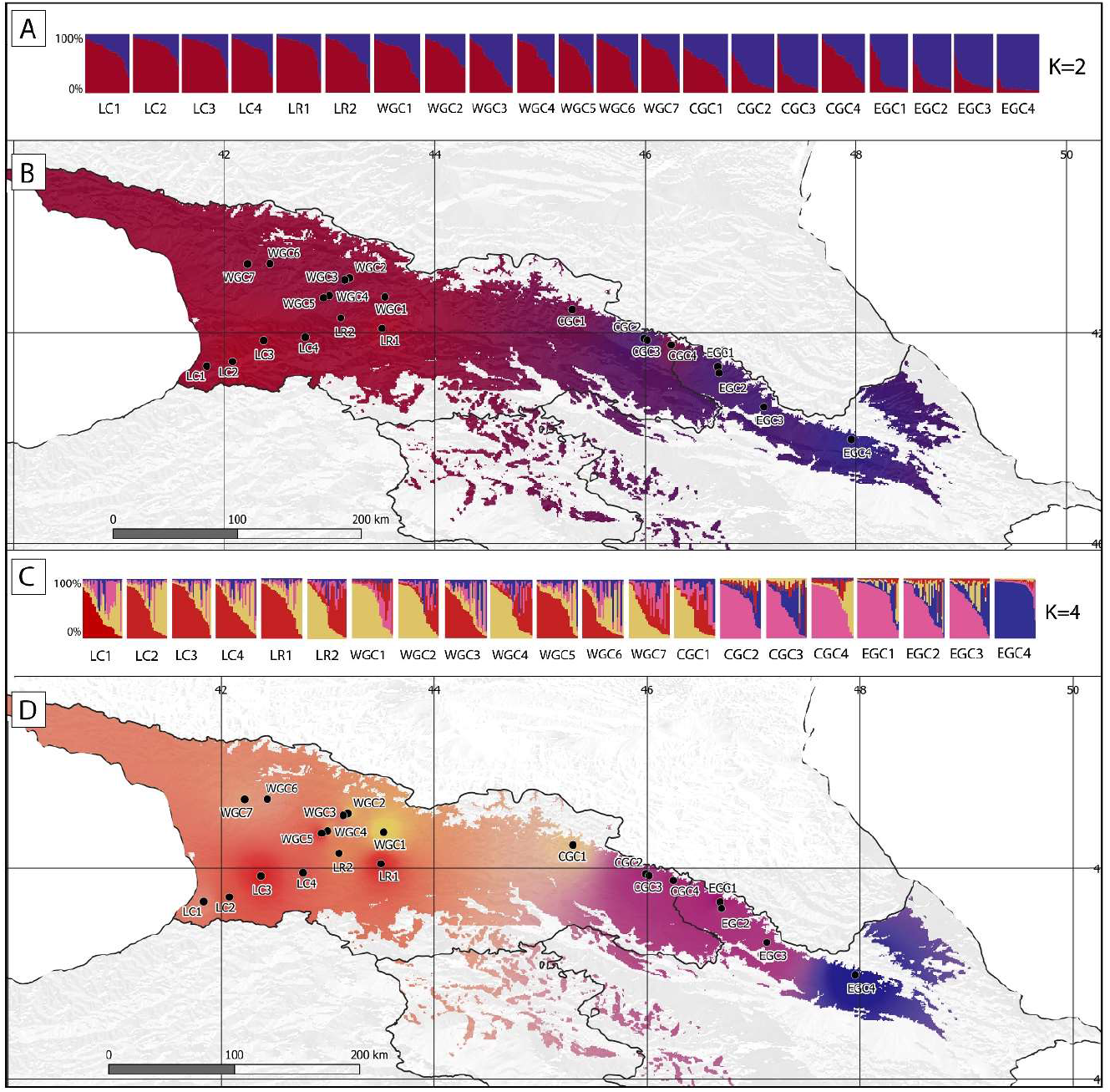
Spatial genetic structure of populations of *C. sativa* in the South Caucasus inferred by STRUCTURE: (A) barplot of individual estimated membership according to an optimal number of clusters K=2; (B) interpolation results throughout the landscape at K=2 based on Q-membership values in QGIS; (C) barplot of individual estimated membership according to a second optimal number of clusters K=4, (D) interpolation results throughout the landscape at K=4 based on Q-membership values in QGIS; population acronym in Table 1.

**Fig. 3.**
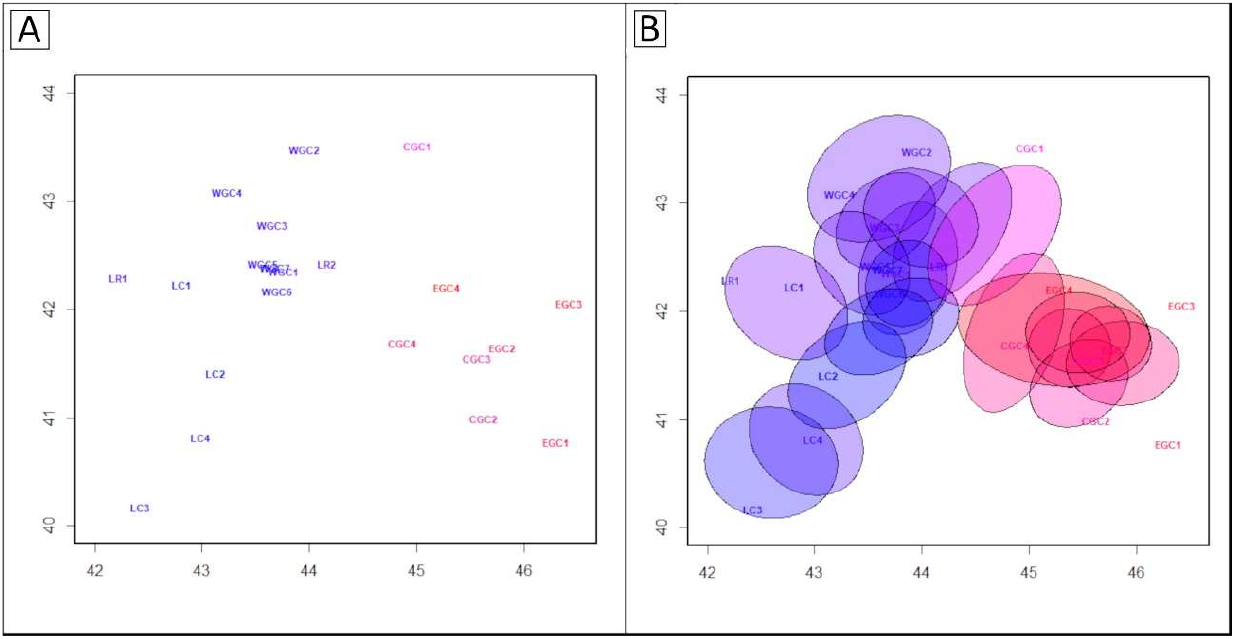
The location of studied populations in geogenetic space using the 50% uncertainty in current geographic locations due to genetic admixture inferred by the SpaceMix analysis. The X and Y axis present longitude and latitude, respectively; populations acronym in Table 1.

We used a 2×10^4^ simulated training data set per scenario for model selection at the first step of ABC-RF analysis. After testing, the number of trees in the constructed random forests was set to 2,000 as this number was large enough to ensure a stable estimation (Supporting information 1, Fig. S1B.). Next, the most likely scenario was evaluated based on a classification vote for each scenario, representing the number of times a scenario is selected in a forest of *n* trees and the posterior probability estimator. The compatibility between scenarios and/or associated priors and the observed data was assessed using the linear discriminant analysis (LDA). After processing the scenario of choice, the second step of ABC-RF analysis included parameter estimation for the most likely scenario. To do this, we performed independent ABC-RF treatments for each parameter of interest based on an extended training data set as 10^5^ simulated data sets, 2,000 trees with 1,000 out-of-bag samples used as a test, five axes of PLS (partial least squares regression analysis) derived from the PLS processed on the summary statistics, and summarised using the 130 statistics that described the genetic diversity within the populations, between pairs or per triples of populations, averaged over all loci used. We inferred the mean and the median as point estimates for each parameter and computed global and local accuracy metrics corresponding to global and local normalised mean absolute error (NMAE) and the 90% coverage.

To evaluate whether our training set is sufficient for ABC-RF analysis, we applied procedures recommended by Collin *et al*. (2021) to compare the accuracy metrics’ stability under different subsets of the training data set. The out-of-bag prediction method (Pudlo *et al*., 2016) based on a 1,000 out-of-bag dataset was used to evaluate the prediction accuracy of the scenario choice and parameter estimations based on global and local error. The global (i.e. prior) error indices correspond to prediction quality measures computed over the entire dataset. In contrast, the local (i.e. posterior) error indices that are conditionally computed on the observed dataset refer to the prediction quality precisely at the position of the observed dataset. For the scenario choice, we conducted ten replicate ABC-RF analyses based on the same training dataset (Collin *et al*., 2021).

In terms of genetic parameters, we used the Stepwise Mutation Model with two parameters: the mean mutation rate (μ) and the mean parameter (P) of the geometric distribution used to model the length of mutation events. We assumed uniform distribution for mean mutation rate and γ for the remaining genetic parameters; their ranges are shown in Table S1 (Supporting information). ABC-RF analysis provides the time from the event (split or merging) in the number of generations that needs to be converted into absolute time. Therefore, the generation time for *C. sativa* was assumed to be 100 years, as recently used by Fernández-López, Fernández-Cruz & Míguez-Soto (2021) in the reconstruction of the demographic history of the species in the Iberian Peninsula to keep the comparability between the results of that study with our investigations.

### Niche modelling and paleobotanical records

The theoretical range of *C. sativa* in the past was estimated to support our demographic estimations and track the location of potential glacial refugia for the species in the Caucasus. For this purpose, we employed the Species Distribution Modelling (SDM) approach based on the maximum entropy algorithm implemented in MAXENT 3.4.3 (Phillips *et al*., 2004) in a series of analyses. Specifically, we aimed to determine the species prevalence in the South Caucasus throughout the following periods: the Last Interglacial (LIG, ca. 140-120 ka), the Last Glacial Maximum (LGM, ca. 21-17 ka), the Early Holocene (EH, 11.7-8.326 ka), the Middle Holocene (MH, 8.326-4.2 ka), and the Late Holocene (LH, 4.2-0.3 ka).

First, our study area was defined within latitudes from 35.77 to 45.16 and longitudes from 38.35 to 56.00. Initially, 214 georeferenced occurrence spots were collected from various sources (Table S1, Supplementary materials 2). However, after data treatment in QGIS to meet the uniform coverage to the study area, in total, 92 unique points were delivered for the further SDMs analysis. Next, a set of bioclimatic variables (Bio1-Bio19) at 30 arc-sec resolution for the current period (1979 – 2013) were downloaded from CHELSA 1.2 (Karger *et al*., 2017). The same set of climatic variables was obtained from the PaleoClim for the past periods, i.e. LIG, LGM, EH, MH, and LH (Fordham *et al*., 2017; Otto-Bliesner *et al*., 2006; Brown *et al*., 2018) with a 2 arc-min resolution. To avoid the multicollinearity among variables in the studied landscape, the *vif* function implemented in the *usdm* R package was used (Naimi *et al*., 2014). Finally, seven variables (Bio1 - Annual Mean Temperature, Bio3 - Isothermality, Bio8 - Mean Temperature of Wettest Quarter, Bio9 - Mean Temperature of Driest Quarter, Bio15 - Precipitation Seasonality, Bio18 - Precipitation of Warmest Quarter, Bio19 - Precipitation of Coldest Quarter) remained for further analyses. MAXENT was run with a “random seed” option, in which 20% of the occurrence was set as test data for model evaluation. Replicates were specified as 100 for bootstrap procedures with maximum iterations ofv10^4^ and output was set as logistic. The area under the curve (AUC) values were used to estimate the performance of each model (Phillips *et al*., 2004; Wang et al., 2007). The high values of AUC (> 0.9) indicate a significant prediction power. SDMs results were visualised in QGIS.

Additionally, to support our SDMs findings, we used published data on pollen and macroremains records from Krebs *et al*. (2019). Specifically, we collected and filtered records to match the Caucasian ecoregion (Supporting information 2, Table S2). Then, we overlaid the occurrence of pollen and macroremains on the maps delivered from MAXENT analysis showing the suitability areas in LGM, EH, MH, and LH using QGIS. In addition, because of the well-known history of cultivation and human influence on the distribution of *C. sativa* in Europe (Conedera *et al*., 2004), it was crucial to divide pollen records into groups that accounted for the probability of human-induced dispersion of the species in the South Caucasus. Therefore, we followed suggestions about the cultivation of the species found in Krebs *et al*. (2019) and divided the collected data for the LH period into pre-cultivation records – (up to 2500 BP) and cultivation records – (after 2500 BP). In total, 89 records were retrived from Krebs *et al*. (2019), of which nine were solely macroremains (Supporting Material 2, Table S1). The pre-cultivation period consisted of 58 records, while a further 31 records were ascribed to the cultivation period (Fig. 5).

**Fig. 4.**
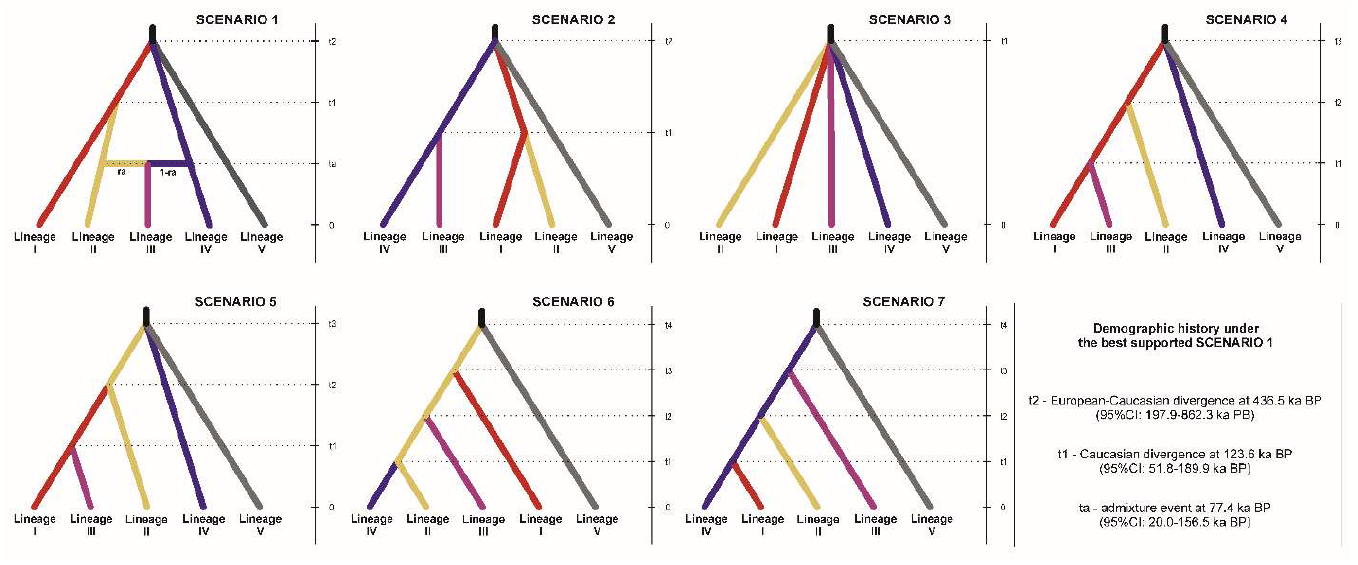
Seven demographic scenarios of the possible evolutionary history of *C. sativa* tested in the ABC analysis. The estimated time of demographic and admixture events were scaled by the species generation time (100 yrs.). Genetic lineages representing the main clusters inferred by STRUCTURE, *i.e*.: Lesser Caucasus (Lineage I, red), Greater Caucasus, (Lineage II, yellow), West+East Greater Caucasus (Lineage III, purple), East Greater Caucasus (Lineage IV, blue) and Europe (Lineage V, grey). According to the computation, Scenario 1 was selected as the most probable one for the species.

**Fig. 5.**
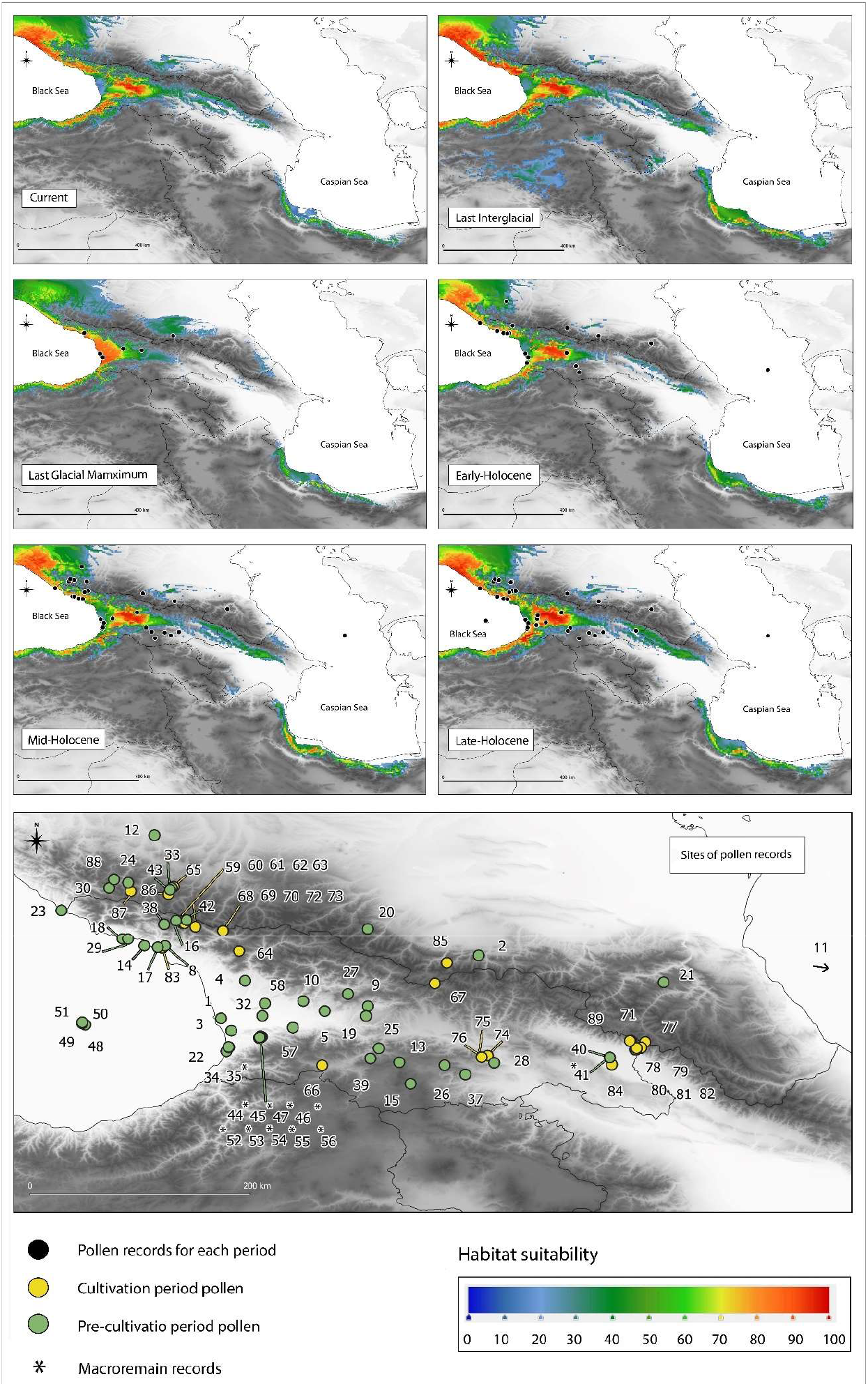
Estimation of the potential range of species’ occurrence in present times (1979 – 2013), the Last Interglacial (LIG, ca. 130 ka), the Last Glacial Maximum (LGM, ca. 21-17 ka), the Early Holocene (EH, 11.7-8.326 ka), the Middle Holocene (MH, 8.326-4.2 ka), and the Late Holocene (LH, 4.2-0.3 ka), with pollen record sites from the corresponding periods according to Krebs *et al*., (2019). Sites of pollen record in *yellow* represent the cultivation period (up to 2500 BP) and *green* the pre-cultivation period (after 2500 BP).

## Results

### Genetic diversity

All analysed nSSR loci were neutral and no linkage disequilibrium between each pair of loci was detected. Null alleles were presented in all loci with a frequency ranging from 0.009 (EMCs13) to 0.120 (CsCAT41) and an average value of 0.004 (Table S1, Supporting information 3). A total of 113 different alleles were found in studied populations ranging from 5 (EMCs2) to 25 (CsCAT41) with a mean value of 12.55 per population.

The genetic structure parameters are presented in Table 1. The highest value of the average number of alleles per locus (*A*) was noted in the two Lesser Caucasus populations (6.67, LC1 and LC4) and the lowest in the Central Greater and Eastern Greater Caucasus (3.56, CGC4 and EGC3) and this pattern was also found in the private alleles frequency where the highest Pa (1.75) was noted in LC populations and the lowest (0.25) in EGC. Observed heterozygosity (*Ho*) ranged from 0.583 (WGC5) to 0.413 (CGC3), while populations from the Lesser Caucasus presented the highest level of *uHe*, reaching a value of 0.594 in LC2, and the lowest was observed for the Central Greater Caucasus stand (0.415 in CGC4). The highest *Fis* was observed in LC4, WCG6, and CGC3 reaching 0.111, 0.131, and 0.136, respectively. The value in the remaining populations was relatively low and remained below 0.1. According to DIC, the homozygosity excess in the populations with the highest *Fis* was likely due to the result of inbreeding.

The single population from the European range represented by Macedonia presented the highest values of private alleles (Table 1) but moderate values of *A* (5.22), *Ar* (4.57), and also highest values of genetic diversity parameters (*Ho*=0.651 and *uHe*=0.652). The inbreeding coefficient was the lowest (*Fis* =0.015) among all studied populations.

The global genetic differentiation among populations was moderate, reaching 0.076 (95% CI: 0.055-0.094), while the *F_ST_* estimation with the ENA correction was slightly lower *F_ST_* =0.073 (95% CI 0.052-0.092). Pairwise Fst ranged from 0.004 (WGC4/WGC3) to 0.266 (EGC4/CGC4). Generally, the most divergent was the most easterly located population, EGC4, which attained the highest values in pairwise comparisons and likely, the distinctiveness of this population affected the overall level of differentiation (Supporting information 3, Table S2).

The regional-level analysis showed statistically significant differences in *Ar* and *He* among the groups of populations in three scenarios out of four tested (Supporting information 3, Table S3). No differences were found in any parameters of the genetic structure between populations from the Lesser Caucasus and West Greater Caucasus. The global differentiation (*F_ST_*) among the groups of populations was significant only when comparing West Caucasian populations (LC, LR, and WGC) to Central and East Greater Caucasian populations (CGC, EGC), reaching the mean values of 0.046/0.095, respectively (*p*<0.0369); no statistical difference was observed in *Fis* values (Supporting information 3, Table S3).

### Range-wide population structure

An optimal number of clusters for the South Caucasus populations and North Macedonian site was K=3 according to the ΔK method, and K=10 in case of LnP(K) (Mean LnP(K) = −13694.3; SD = 12.63179), while the approach of Puechmaille (2016) retrieved five clusters from the data (Supporting information 3, Figs. S1-S5). Both clustering approaches underpinned the outstanding genetic composition of the Macedonian population. Furthermore, the clustering at K=5 delivered an exact differentiation pattern as an analysis conducted for the South Caucasus subset of populations described below (Fig. 1, Supporting information 3, Fig. S5).

Based on LnP(K), the optimal number of homogenous groups in the Caucasian dataset was K=10 (Mean LnP(K) = −13185.36; SD = 140.42), which was too high to provide any other explanation, but the idiosyncratic history of each population (Supporting information 3, Fig. S6). In contrary, Evanno’s *ΔK* method applied to the Caucasian populations (Georgia and Azerbaijan) indicated that at K=2, the data was the most optimally structured (Supporting information 3, Fig. S7). This grouping pattern assumed Western (Cluster I: populations WGC and LC) and Eastern (Cluster II: populations CGC and EGC) gene pools and clearly depicted admixture between them (Fig. 2 A and B). The population CGC1 located just between those two gene pools exemplified this genetic admixture as it displayed almost equal average membership to both clusters (I: 52% and II: 48%). However, an outlier from the general pattern was the CGC4 population, with a high proportion of Cluster I (56%) being located in the region of the predominance of Cluster II.

Further details in the genetic structure were delivered by using K=4, based on the method of Puechmaille (2016) (Fig. 2C and D, Supporting information 3, Fig. S8). Generally, K=4 indicates sub-structuring in the western (the Colchis) and eastern part of the species range, also revealing a more complex pattern of genetic structure. Specifically, the population from the Lesser Caucasus (LC1-LC4) were grouped in Cluster I together with two populations from the West Greater Caucasus (WGC3 and WGC5) and a population from the eastern edge of the Likhi Range (LR1), while Custer II had wider spatial distribution and consisted of three populations from the West Greater Caucasus (WGC1, WGC2, WGC4), LH2 from the western edge of the Likhi Range and single population from the Central Greater Caucasus (CGC1). However, the majority of populations in both mountain ranges display a quite high admixture with no predominance of any of the two clusters that have been defined. This is well exemplified in WGC6, WGC7 and CGC1, which possessed almost equal membership levels (Q) to Cluster I and II. The highest Q of Cluster I had LC3 (61.8%), while Cluster II - WGC1 (61.4%). Cluster III was defined for populations distributed in the Central Greater Caucasus (CGC2-CGC4) and the East Greater Caucasus (EGC1-EGC3). Here, we also noted some fraction of the genomes (ca. 20%) attributed to the cluster I and II. At the easternmost part of the studied area, the population EGC4 of Cluster IV shows a high divergence to the other populations studied, confirmed by the highest values of pairwise *Fst* reaching >10%. The admixture of this distant genetic pool at levels of 24.5% and 22.4% were detected in EGC3 and EGC2, respectively.

The constructed geogenetic map made of population relationships by SpaceMix is given in Figure 3. Plots of the posterior probability trace and model adequacy indicated that MCMC mixed well and achieved the acceptance rate close to the desired 44%, and the model adequately described the data (Supporting information 3, Fig. S9A, B). Basically, the SpaceMix analysis recovered the spatial pattern of two major groups that have been revealed by STRUCTURE’s K=2 with the intermediate position occupied by CGC1 (Fig. 3A). The picture obtained via SpaceMix primarily shows the west-east spatial dimension of the divergence, but it also suggests a north-south pattern that reflects some distinctiveness between the Greater and Lesser Caucasus ranges. The plotted populations’ uncertainty in location shows that populations are generally located in one of the two groups. However, for some populations, their geogenetic locations are significantly pulled away from their current geographic one (e.g., CGC1, CGC2, EGC1, and EGC3), which likely reflects genetic relatedness with other populations and long-distance admixture (Fig. 3B). On the other hand, their geogenetic locations are closer for some populations despite the considerable geographic distance. The detailed insight into the admixture pattern reveals persuasive gene flow and variability of admixture in terms of direction. For example, population CGC1 reflects substantial admixture from the geogenic space related to the Lesser Caucasus, while LR2 to the Greater Caucasus (Fig. 3B).

### Demographic history

The classification votes and posterior probabilities estimated for the observed microsatellite dataset in all ten replications of ABC-FR analyses were the highest for Scenario 1 (average of 646 (SD=21.417) votes out of 2,000 RF-trees; average posterior probability PP=0.722). The global probability of incorrectly identified scenarios (prior error rates) in tested datasets was estimated to be 0.242 (Supporting information 1, Table S2) and the highest-class error rates were in Scenarios 2 and 7 (0.423 and 0.424, respectively). For Scenario 1, the posterior (local) error rate was on average 0.278. The fit of Scenario 1 to the observed dataset is illustrated by the projection of the simulated and observed datasets on the first two LDA axes (Supporting information 1, Fig.S1).

Under the most likely scenario (Fig. 4) at *t2*=436.5 ka BP (95% CI: 197.9 - 862.3 ka BP), the ancestor lineage split into Lineages I, IV, and V which is primarily translated into a disjunction between the Caucasian and European gene pool. Additionally, this scenario supported the high genetic distinctiveness of Lineage IV originating from ECG4. At *t1*=123.6 ka BP (95% CI: 51.8-189.9 ka BP), Lineage II, having the highest frequency in West Greater Caucasus, diverged from Lineage I that predominates in the Lesser Caucasus. Finally, the almost symmetrical admixture between Lineage II and IV dated at *ta*=77.4 ka BP (95% CI: 20.0-156.5 ka BP) led to Lineage III. The highest effective population size was estimated for Lineage I, and the lowest for Lineage III (Table 2).

**Table 2.**
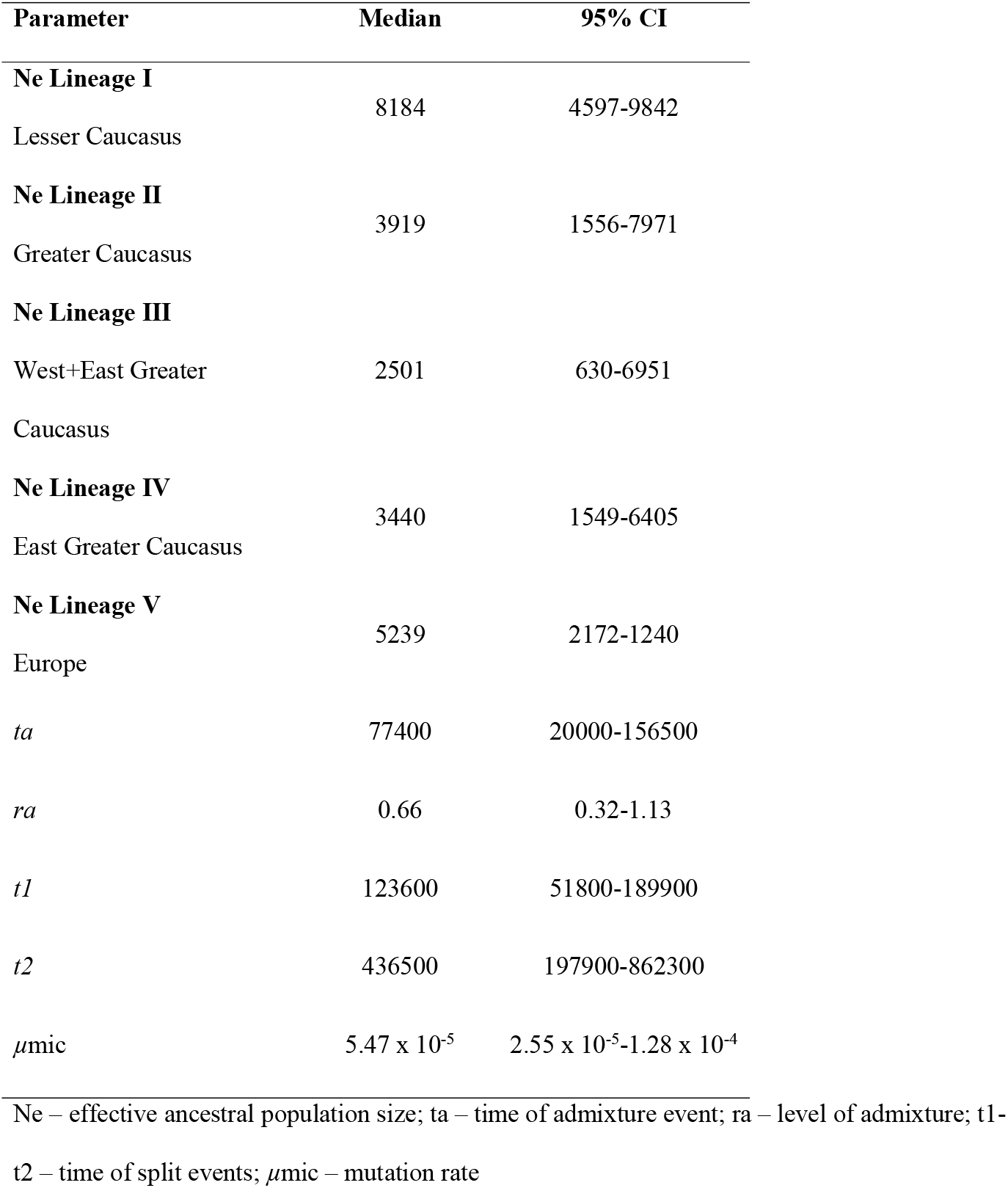
Parameter estimates for the best demographic Scenario 1 performed in DIYABC Random Forest

### Niche modelling and paleobotanical records

The MAXENT model accuracy (AUC) for all presented SDM models was greater than 0.93 (Supporting information 2, Table S2). Depending on the model, the most important variables in the SDMs were the precipitation of the warmest quarter (Bio18) and the precipitation of the coldest quarter (Bio19), with a relative contribution reaching >54% and >21%, respectively. In addition, the annual mean temperature (Bio1), with a relative contribution of >11%, also had a significant impact on the projected species distribution in all models. Detailed information about the contributions of all bioclimatic variables is given in Table S2 (Supporting material 2).

The potential distribution of *C. sativa* across the studied landscape in different time slices is given in Figure 5. The current theoretical range overlaps with the known current species distribution. In terms of niche suitability, the most suitable habitats were in the westernmost part in the foothills of the Western Greater Caucasus, Abkhazia and East Pontic Mts. in Turkey. Towards the east, habitat suitability declines, reaching a maximum value of 20-40% in the Central Greater Caucasus and East Greater Caucasus (Azerbaijan). The second domain of the potential distribution covers Hyrcanian forests in Iran, where the current species presence is limited to a few stands only in the westernmost part of the region. However, higher potential suitability of 50-70% in comparison to the eastern part of the Greater Caucasus was projected for sweet chestnuts in the Hyrcanian forest.

The niche model for LIG indicates that the distribution of the species covered approximately the same areas as the current range. Additionally, MAXENT modelled small pockets of the species occurrence in the Azerbaijan part of the Greater Caucasus (suitability up to 60%), the Lesser Caucasus (up to 40%), central Armenia and southern territories of Turkey where the species is not present currently. The projected distribution in Hyrcania in LIG is wider than currently with much higher suitability. There was a clear distributional gap between Hyrcanian forests and the Greater Caucasus in that period.

The modelled distribution contracted at the LGM. Chestnut probably found suitable climatic conditions mostly in Colchis Lowland near the Black Sea and a narrow strip of coastal areas close to the Pontic Mts. The Hyrcanian forest projected a relatively high suitability of conditions in LGM for *C. sativa* (40-60%) and the North Caucasus (Kabardino-Balkaria-North Ossetia) as well (suitability ca. 50%). The model for the early Holocene (EH) suggests an expansion of the distribution to the east compared to LGM and possibly an expansion in the Hyrcanian forest. In the Mid-Holocene (MH), the range remained more or less the same in terms of the area, with increased suitability in eastern locations. The only noticeable difference by the Late Holocene (LH) was lower suitability values in the Hyrcanian forest compared to MH, but still higher than it is today.

The compiled fossil pollen records retrieved from Krebs et al. (2009) are largely in line with the modelled species range from the LGM to the current times in terms of the projected distribution in each analysed period (Fig. 5). The data support the modelled presence of *Castanea* during LGM in areas indicated with moderate (50%) and high (90-100%) climatic suitability. For EH, the fossil pollen of sweet chestnut was reported in cores taken from sites beyond the projected range, i.e. in the Javakheti Plateau and in some locations in the Northern Caucasus, undoubtedly a result of long-distance pollen transport. Similarly, the cores from the Caspian Sea contained sweet chestnut pollen likely refers to the wind transportation of pollen grains from the Hyrcanian forests.

## Discussion

### The European-Caucasian divergence

Our modelling suggests that the genetic split between the European and the Caucasian gene pools of *C. sativa* occurred during the Middle Pleistocene (465.5 ka, 95% CI: 197.9 - 862.3 ka). This period marks the end of the pronounced shift in the cyclicity of Quaternary glaciations called the Early Middle Pleistocene Transition (EMPT) that spanned the period of 1.2 - 0.6 Ma (Head & Gibbard, 2015; Chalk *et al*., 2017). The weaker and shorter glacial cycles with a ~40 000 yrs frequency that predominated in the first ca. 1.5 million yrs of the Pleistocene gave way to longer and stronger oscillations lasting 100,000 yrs. Consequently, the glaciations that took place after EMPT were generally longer, colder and drier and led to the formation of larger continental ice sheets and marked vegetation turnover (Chalk *et al*., 2017). In the West Caucasus, a reconstruction of Miocene to Pleistocene climate conditions indicated decreasing temperatures since the beginning of the Pleistocene with a significant drop in temperatures at the beginning of the Middle Pleistocene (0.85 Ma), which was accompanied by a shift in the vegetation from a more warm-temperate character to its current type (Mahler, Shatilova & Bruch, 2022).

It seems very likely that the European-Caucasian genetic divergence for *C. sativa* was driven by the generally cooler and dryer climate after the EMTP culminating in the severe glacial/interglacial cycles of the Middle Pleistocene. This development led to a climatic differentiation between Europe and the Caucasus region as the latter was additionally strongly affected by orogenic uplift. The rise of the Greater Caucasus and the Hyrcanian mountains protected the eastern coast of the Black Sea as well as the southern shores of the Caspian Sea from aridification and supported the development of humid refugia (Mahler, Shatilova & Bruch, 2022). In Europe, a stepwise decline of arboreal pollen is observed during the Middle Pleistocene. For example, on the Balkan Peninsula, the glacial Marine Isotope 22 (MIS 22) just before the beginning of the Middle Pleistocene led to a major vegetation turnover, while the following glacial maxima during the Middle Pleistocene (mainly MIS 16 and MIS 12) contributed to the final reduction of *Castanea* that later has never been an abundant component of the interglacial forests of Europe (Tzedakis *et al*., 2006). Also, the Pyrenees lineages of *Abies alba* Mill. diverged from its ancestral Alpine lineages during the EMPT (Scotti-Saintagne *et al*., 2021). In the Apennine Peninsula, mesophilous tree species such as *Juglans* responded with a range reduction for EMPT (Orain *et al*., 2015). Further progressive disappearances of several tree species as a response to EMPT are reported from pollen records from North Spain, South France and Central Italy (Magri & Palombo, 2013).

In addition to the cooling trend, increased summer aridity had a severe impact on vegetation change, documented for the Mediterranean realm (Postigo-Mijarra *et al*., 2010; Magri & Polombo, 2013), and also shown in our data by the high relevance of the precipitation of the warmest quarter (Bio18) for the MAXENT’s models. However, precipitation remained relatively stable in the West Caucasus since the Neogene with less humid conditions than today (Mahler, Shatilova & Bruch, 2022). Rather than being driven by global climatic changes, the Pleistocene evolution of precipitation in the West Caucasus can be attributed to alpine orogenic uplift. The increasing altitudes of the Greater Caucasus and especially the Likhi Range during the Quaternary (Sosson *et al*., 2010) had a significant influence not only as biogeographic barriers but also caused a stepwise reorganization of the precipitation regime in the South Caucasus. While precipitation decreased considerably in Europe from Pliocene onwards (Mosbrugger, Utescher & Dilcher, 2005), humidity remained high in the South Caucasus (also during summer) or even slightly increased in the West Caucasus due to the rain shadow effect of the rising Likhi Range (Mahler, Shatilova & Bruch, 2022). These changes in humidity, evolving through the interplay of global climate change and tectonic processes, probably were the most influential driving force for the reorganisation of the Quaternary vegetation and the split of forest distributions in the Caucasus. To some extent, the region’s climate followed the global trend of temperature decrease known as the Late Miocene Cooling that started at ca. 14 Ma. While in Europe, it was accompanied by a pronounced drop in precipitation (Mosbrugger, Utescher & Dilcher, 2005), this was not observed in western Georgia (Mahler, Shatilova & Bruch, 2022). The precipitation in the western territories stayed high compared to the eastern part of the Caucasus because of the buffering effect of the mountain barriers (the Greater and Lesser Caucasus and Likhi Range) blocking the moisture supplied by westerly airstreams originating over the Black Sea (Tarkhnishvili, 2014). Today, the vegetation of the Caucasus is organised along the precipitation gradient formed due to orogenic activity in the region – the western territories are covered with moisture-demanding biomes, and the eastern parts are inhabited by arid and semi-arid plant communities (Connor & Kvavadze, 2009). The natural ranges of *Picea orientalis* ((L.) Peterm.) and *Abies nordmanniana* ((Steven) Spach.) which concentrate in western and central Georgia overlap with the distribution of the precipitation in the Caucasus and haplotypic diversity in *Q. petaea* subsp. *iberica* reflects the impact of isolation due to the geographic barrier of Likhi Range (Ekhvaia *et al*., 2018). Recent work of Sękiewicz *et al*. (2022) suggest that the genetic diversity and differentiation patterns in *Fagus orientalis* (Lipsky) were also shaped by climatic distinctiveness between western and eastern regions of the Caucasus.

### The Middle to Late Pleistocene genetic splits and admixture in the Caucasus

The strong signal of east-west divergence in the sweet chestnut Caucasian gene pool was confirmed by two different analyses of our dataset. First, the Bayesian cluster analysis revealed two clusters that captured the major structure in the data. The SpaceMix approach repeated this east-west split in genetic structure. A similar genetic pattern has been found for two other relic species, *Zelkova carpinifolia* (Maharramova *et al*., 2015) and *Pterocarya fraxinifolia* (Christie *et al*., 2014), denoting the Colchic-Hyrcanian divergence and the role of both subregions as refugia during LGM.

A much deeper structure was revealed when the second optimal number of clusters, K=4, was applied to the data. This showed a clear distinctiveness of the marginal population ECG4 at the eastern extreme of the range, belonging to its own Cluster IV. There are currently some remnant populations of the sweet chestnut still growing in the easternmost part of the range in Hyrcania (Iran) that are under high conservation risk due to low diversity, fragmentation and overexploitation (Janfaza *et al*., 2017). Between the areas of sweet chestnut occurrence in the East Greater Caucasus and Hyrcania, there is a large distributional gap in forest vegetation suggested by the MAXENT’s models, at least since LIG. This is due to a vast semi-arid area in the driest part of the South Caucasus, covered by semi-deserts and xeric shrublands (Peper, Pietzsch & Manthey, 2010). According to the most optimal demographic scenario, the genetic split between Lineage IV located in Azerbaijan and originating from ECG4 and Lineage I in the Lesser Caucasus was as ancient as the European-Caucasian divergence, i.e. in the Middle Pleistocene. Accordingly, the first diversification in the South Caucasian gene pool of the *C. sativa* was probably induced by the same drivers. The Pleistocene uplift of the Likhi Range and its rain shadow effects that were the reason for stable humid conditions in the West Caucasus, at the same time caused a considerable decrease in precipitation in the eastern lowlands of the Caucasus and thus a severe east-west gradient in humidity separating forest populations (Shatilova *et al*., 2011; Mahler, Shatilova & Bruch, 2022).

A current lack of samples from the Hyrcanian populations of *C. sativa* makes it difficult to settle the genetic relationships of ECG4 to Hyrcanian populations despite the prominent difference of ECG4. Similarly, eastern Georgian populations of *Zelkova carpinifolia* were reported to carry haplotypes of the Hyrcanian group despite obvious geographical proximity to the Colchis (Christie *et al*., 2014), which supports the idea that ECG4 may originate from the Hyrcanian refugium, or at least, has strong genetic ties to that region. The genetic distinctiveness of the lineage present in ECG4 might be acknowledged to adaptation processes which have been documented to produce spatial genetic patterns in trees (Mayol *et al*., 2010; Orsini *et al*., 2013). Recently, adaptation to rainfall has been reported in European populations of sweet chestnut growing in dry regions of Spain, Greece and Italy, which designates the species’ potential to respond to this selection factor (Alcaide *et al*., 2019; Castellana *et al*., 2021).

The inferred timing of the split between Lineage I and Lineage II (123.6 ka BP; 95% CI: 51.8-189.9 ka BP) overlaps with the second part of the Middle Pleistocene and represents the penultimate glacial/interglacial cycle of MIS 6 and MIS 5 (Lisiecki & Raymo, 2005; Krijgsman *et al*., 2019). Palynological data retrieved from the Southern Black Sea coast indicated the occurrences of pollen of *Castanea* mainly during the LIG (MIS 5e) (Shatilova *et al*., 2011), while during the preceding glacial *Castanea* pollen emerged only sporadically during times of slightly increased temperature as indicated by melt water signals, and probably related to increased humidity (Shumilovskikh *et al*., 2013). However, assuming a separation of populations necessary for a genetic split, a divergence during the preceding glacial MIS 6 seems to be more likely than at MIS 5e, and falls within the confidence interval above. This severe glacial period has been recently shown to also play a key role in the evolution and distributional pattern of the herbaceous *Bunias orientalis* L. in Europe and the Caucasus (Koch *et al*., 2017).

The last identified event in the evolutionary history of sweet chestnut in the Caucasus involved the secondary encounter between Lineage IV and Lineage II. It is dated as 77.4 ka BP (95% CI: 20.0-156.5 ka BP), which roughly coincides with the beginning of the last glacial period (Krijgsman *et al*., 2019). The admixture between those lineages was nearly symmetrical and gave rise to Lineage III, distributed currently in East Greater Caucasus (populations CGC2-EGC3). The confidence interval locates the probable meeting from as recently as the LGM to as early as the LIG. The interglacial period seems to be more favouring for admixture event. After LIG, the Northern Hemispheres experienced climate deterioration and advancement of the glacial sheets that culminated in LGM (Shackleton *et al*., 2021). The reduction of *C. sativa* occurrence in the region was suggested by results of modelling for that period. Paleoclimate reconstruction for West Georgia showed significant cooling and the lowest temperatures, although warm climate taxa are still detected in pollen spectra from that time. At the same time, in East Georgia, the vegetation was reduced and glacial steppe species predominated (Shatilova *et al*., 2011).

Therefore, we argue that the admixture detected in our study is less likely during glacial period as indicated by demographic reconstructions. Conversely, considering that the confidence interval includes the LIG, the admixture may have more likely happened due to forest expansion during this favourable period of increased temperatures and precipitation. The vegetation reconstruction for West Georgia based on pollen data indicates warm and humid conditions at that time and prominent occurrences of deciduous species (Mahler, Shatilova & Bruch, 2022), including fossil leaves of *C. sativa* (Shatilova *et al*., 2011). Range modelling performed here for *C. sativa* for LIG suggest approximately the same areas as the current range with even wider occurrence at the eastern margins in the Greater Caucasus. Hence, the precise time frame and drivers of admixture between Lineages II and IV remain an open question in our work. To resolve this, our study should to be supplemented with wider and more detailed sampling in the eastern part of the region (Azerbaijan), the Hyrcania and also isolated stand in Armenia.

### Refugia during the last glacial cycle and current diversity patterns

Accurately assessing areas of species refugia should integrate different sources of information since approaches based on fossils, genetic data or SDMs are each subject to limitations and biases. In the Caucasus, pollen data recorded vegetation changes since LGM with quite satisfactory resolution and indicated the location of major refugia in Colchis and Hyrcania (Connor, 2006; Messager *et al*., 2021), which finds support in SDMs and genetic data (Tarkhnishvili *et al*., 2012; Aradhya *et al*., 2017; Parvizi *et al*., 2019; Volkova *et al*., 2020; Dering *et al*., 2021). According to SDM analysis, *C. sativa* could have survived LGM mostly in the Colchis refugium and partly in the eastern Pontic Mts. However, the Hyrcanian forests were also suitable and likely refugia. Generally, glacial refugia are expected to harbour higher levels of genetic variability compared to areas that have been recolonized after the LGM. This is especially true for allelic richness and private alleles (Hewitt, 2000). In our study, this was found in populations from the western Georgian Lesser Caucasus (LC1-LC4), which display the highest gene diversity and number of private alleles. Additionally, the genetic lineage that originates from this area (Lineage I) was estimated to have the largest ancestral effective population size, which undoubtedly played a vital role in buffering the erosion of diversity and providing resilience. The niche modelling also suggests that sweet chestnut survival was feasible in a limited area of the easternmost part of the Greater Caucasus, i.e. in the East Greater Caucasus in Azerbaijan (Fig. 5). In paleobotanical records from the Khunzakh plateau (southern Dagestan), *C. sativa* was reported from Early Holocene deposits (9200 to 8980 cal BP; Ryabogina *et al*., 2019), which may indicate the local presence of the species and its spread from the Eastern Greater Caucasus refugial area. However, examples from other species might shed more light on this region and its likely function as a last glacial refugium. In this way, another important refugium previously not considered for the Caucasus might be added to the two major ones already widely acknowledged.

After LGM, according to paleoclimate reconstructions, forest expansion was delayed in the region due to limited humidity during the growing season despite an overall increase in annual precipitation (Messager *et al*., 2021). The Early Holocene model showed a wide range of *C. sativa*, almost comparable to current times. However, the most eastern part of the range was likely to be unsuitable and the colonisation could be hampered due to relatively dry climate in eastern Georgia at the beginning of the Holocene. The stands currently located in this areas display the lowest values of the gene diversity. The bottleneck effect during colonisation of the eastern areas and exposition to drift in much more challenging environment might be a reason of observed genetic impoverishment as recently argued for other species in the region (Ekhavaia *et al*., 2018). While EH is considered to be drier than today, at least during growing season, the MH climatic optimum should have been a very suitable time for sweet chestnuts, which has been supported by the MAXENT projection. The abundance of *Castanea* pollen in cores dated to MH has been documented for western Georgia (Kavavdze & Connor, 2005). Late Holocene is the time of the almost final formation of the range of sweet chestnut, which currently covers West Georgia and Abkhazia, supported by pollen data of Connor, Thomas & Kavavdze (2007).

Despite the rugged landscape of the Caucasus, the STRUCTURE and SpaceMix both revealed intensive gene flow among populations of sweet chestnut as it was found earlier for other tree species in the region (Maharramova *et al*., 2015; Dering *et al*., 2021). Both methods showed the prevailing gene exchange between populations located in western Georgia, i.e. between the Lesser and Greater Caucasus. The moderate gene flow was indicated among the populations from Central and East Greater Caucasus. This may point to the Likhi Range as a significant biogeographic barrier preventing wide genetic connectivity which was detected also for *Q. petrea* subsp. *iberica* (Ekhaia *et al*., 2018). Nevertheless, the gene flow remains quite intensive, ascribed to the relatively small dimensions of the studied area and the potential of long-distance dispersal due to the small size of the pollen grains and the high rate of pollen production (Larue *et al*., 2021). Although predominantly insect-pollinated, sweet chestnut may use wind as a dispersal agent, as found in another Neogene relic tree species, *Aesculus hippocastanum* L. (Thomas *et al*., 2019) resulting in intensive gene flow noted for this species in natural range (Walas *et al*., 2019). The homogenising effect of gene flow among populations of the sweet chestnut in the Caucasus is reflected in a moderate level of differentiation, reaching 7%. The level of admixture on a pan-European scale definitely remains much more limited, which has been confirmed by a clear geographic structure and a relatively high level of differentiation reaching 15% (Mattioni *et al*., 2013; Castellana *et al*., 2021). Additionally, the results of SpaceMix underlined the complexity of the coalescence process involving inherent temporal and spatial aspects (Bradburd & Ralph, 2019). The geographic closeness of the populations does not imply their genetic relatedness and *vice versa* because of modelling impact of the gene flow on genetic composition, especially long-distance one. For that reason, some uncertainty was inferred between the current locations of populations and their genetic composition and source of admixture. This is evident in the easternmost part of the range of *C. sativa*, which may include the hypothetical impact of the Hyrcanian gene pool.

### Final remarks

In contrast to the European gene pool of sweet chestnut that has been substantially altered by cultivation and propagation (Fineschi *et al*., 2000; Martín *et al*., 2012; Fernández-López, Fernández-Cruz & Míguez-Soto, 2021) the utilisation of the sweet chestnut in the Caucasus was limited to harvesting timber, nuts, and fodder and honey production but without large-scale commercial planting which was restricted to small orchards using local seeds (Bobokashvili & Maghradze, 2009). Moreover, severe and uncontrolled cutting of the tree occurred in Georgia during the 19th Century, due to high European demands for sweet chestnut timber, had a pronounced negative impact on the species, leading to the highly fragmented distribution seen today (Bobokashvili & Maghradze, 2009). During the Soviet period, attempts to cultivate the species were undertaken in the North Caucasus (Abkhazia, Russia), but mostly failed due to fungal and bacterial blight outbreaks (Yazvenko, 1994; Pridnya, Cherpakov & Paillet, 1996). Hence, the biogeographic patterns uncovered for sweet chestnut in the Caucasus are likely less disturbed than in Europe and more likely to reflect natural evolutionary changes. This is partially supported by fossil pollen data (Krebs *et al*., 2019). There is no detectable increase of sites with *Castanea* pollen after 2500 BP which could suggest a spread of the species due to human activity (see Fig. 5).

Most of today’s populations of sweet chestnut are seriously fragmented and devastated by *Cryphonectria parasitica* leading to a mass die-off of trees, especially in Georgia (Dumbadze *et al*., 2018; Tavadze *et al*., 2013). Due to sweet chestnut’s longevity, the adverse effects of this are not yet detectable – populations display generally high gene diversity and low inbreeding. However, the widespread scale of *C. parasitica* infections in Georgia and its dynamic expansion toward Azerbaijan since 2003 (Aghayeva & Harrington, 2008) is alarming and puts the populations at high risk of future decline. Although the regional impact of future climate change in the Caucasus is not yet fully understood, especially regarding precipitation changes (IPCC, 2014, Figure SPM.7), it will likely amplify the biotic stress on sweet chestnut. The resulting habitat changes and shifts in the range of species will have further negative impacts on biodiversity hotspots (IPCC, 2021), including probably this one. As yet, sweet chestnut, as a relict tree species in the Caucasus, is not under the conservation targets. This urgently needs rectifying.

## Acknowledgements

This work was supported by the National Science Centre [NCN, Project Number UMO-2017/26/E/NZ8/01049], the Institute of Dendrology, Polish Academy of Sciences and Poznań University of Life Sciences. We are highly grateful to Georgian and Azerbaijan foresters who supported us during field expeditions in those countries. We also thank M. Łuczak for lab work and G. Iszkuło for assistance during fieldwork.

## References

Adamia S, Zakariadze G, Chkhotua T, Sadradze N, Tsereteli N, Chabukiani A, Gventsadze A. 2011. Geology of the Caucasus: a review. Turkish Journal of Earth Sciences 20(5): 489–544.

Aghayeva DN, Harrington TC. 2008. First report of *Cryphonectria parasitica* on chestnut (*Castanea sativa*) in Azerbaijan. Plant Pathology 57(2): 383–383

Alcaide F, Solla A, Mattioni C, Castellana S, Ángela Martín M. 2019. Adaptive diversity and drought tolerance in *Castanea sativa* assessed through EST-SSR genic markers. Forestry: An International Journal of Forest Research 92: 287–296.

Aradhya M, Velasco D, Ibrahimov Z, Toktoraliev B, Maghradze D, Musayev M, Bobokashvili Z, Preece JE. 2017. Genetic and ecological insights into glacial refugia of walnut (*Juglans regia* L.). PLoS One 12: e0185974.

Arpe K, Leroy SAG, Mikolajewicz U. 2011. A comparison of climate simulations for the last glacial maximum with three different versions of the ECHAM model and implications for summer-green tree refugia. Climate of the Past 7(1): 91–114.

Ballog RA, Maloy RE. 1981. Neogene palynology from the southern California continental borderland, site 467 In: Yeats RS et al. 1981. Initial Reports of the Deep Sea Drilling 63: 565–576.

Barrón E, Rivas-Carballo R, Postigo-Mijarra JM, Alcalde-Olivares C, Vieira M, Castro L, Pais J, Valle-Hernández M. 2010. The Cenozoic vegetation of the Iberian Peninsula: a synthesis. Review of palaeobotany and palynology 162(3): 382–402.

Bobokashvili Z & Maghradze M. 2009. Georgia. In: Avanzato D. (ed). Following chestnut footprints (Castanea spp.): cultivation and culture, folkrore and history, traditions and uses. Scripta Horticulturae 9: 48–52.

Boroń P, Wróblewska A, Binkiewicz B, Mitka J. 2020. Phylogeny of Aconitum subgenus Aconitum in Europe. Acta Societatis Botanicorum Poloniae: 89(3).

Bradburd GS, Ralph PL. 2019. Spatial Population Genetics: It’s About Time. Annual Review of Ecology, Evolution and Systematics 50: 427–449.

Bradburd GS, Ralph PL, Coop GM. 2016. A Spatial Framework for Understanding Population Structure and Admixture. PLoS genetics 12: e1005703.

Brown JL, Hill DJ, Dolan AM, Carnaval AC, Haywood AM. 2018. PaleoClim, high spatial resolution paleoclimate surfaces for global land areas. Scientific Data 5:180254.

Bruch A, Utescher T, Mosbrugger V, 2011. Precipitation patterns in the Miocene of Central Europe and the development of continentality. Palaeogeography, Palaeoclimatology, Palaeoecology 304: 202–211.

Buck EJ, Hadonou M, James CJ, Blakesley D, Russell K. 2003. Isolation and characterization of polymorphic microsatellites in European chestnut (*Castanea sativa* Mill.). Molecular Ecology Notes 3(2): 239–241.

Castellana S, Martin MÁ, Solla A, Alcaide F, Villani F, Cherubini M, Neale D, Mattioni C. 2021. Signatures of local adaptation to climate in natural populations of sweet chestnut (*Castanea sativa* Mill.) from southern Europe. Annals of Forest Science 78(2): 1–21.

Caudullo G, Welk E, San-Miguel-Ayanz J. 2017. Chorological maps for the main European woody species. Data in Brief 12: 662–666.

Chalk TB, Hain MP, Foster GL, Rohling EJ, Sexton PF, Badger MPS, Cherry SG, Hasenfratz AP, Haug GH, Jaccard SL, Martínez-García A, Pälike H, Pancost RD, Wilson PA. 2017. Causes of ice age intensification across the Mid-Pleistocene Transition. Proceedings of the National Academy of Sciences of the United States of America 114: 13114–13119.

Chapuis MP, Estoup A. 2007. Microsatellite null alleles and estimation of population differentiation. Molecular Biology and Evolution 24(3): 621–631.

Christe C, Kozlowski G, Frey D, Bétrisey S, Maharramova E, Garfì G, Pirintsos S, Naciri Y. 2014. Footprints of past intensive diversification and structuring in the genus Zelkova (Ulmaceae) in south-western Eurasia. Journal of Biogeography 41(6): 1081–1093.

Chybicki IJ. Raw2Gen software. Available at:https://www.ukw.edu.pl/pracownicy/strona/igor_chybicki/software_ukw/

Chybicki IJ, Burczyk J. 2009. Simultaneous estimation of null alleles and inbreeding coefficients. Journal of Heredity 100: 106–113.

Collin FD, Durif G, Raynal L, Lombaert E, Gautier M, Vitalis R, Marin JM, Estoup A. 2021 Extending approximate Bayesian computation with supervised machine learning to infer demographic history from genetic polymorphisms using DIYABC Random Forest. Molecular Ecology Resources 21(8): 2598–2613.

Conedera M, Krebs P, Tinner W, Pradella M, Torriani D. 2004. The cultivation of *Castanea sativa* (Mill.) in Europe, from its origin to its diffusion on a continental scale. Vegetation History and Archaeobotany 13(3): 161–179.

Conedera M, Tinner W, Krebs P, de Rigo D, Caudullo G. 2016. *Castanea sativa* in Europe: distribution, habitat, usage and threats. European Atlas of Forest Tree Species 78–79.

Conedera M, Krebs P, Gehring E, Wunder J, Hülsmann L, Abegg M, Maringer J. 2021. How future-proof is Sweet chestnut (*Castanea sativa*) in a global change context? Forest Ecology and Management 494: 119320.

Connor SE. 2006. A promethean legacy: late quaternary vegetation history of southern Georgia, Caucasus (Doctoral dissertation available at: https://minerva-access.unimelb.edu.au/items/a199a5b2-86b5-50b3-9384-3320f9fbdc54)

Connor SE & Kvavadze EV. 2009. Modelling late Quaternary changes in plant distribution, vegetation and climate using pollen data from Georgia, Caucasus. Journal of Biogeography 36(3): 529–545.

Cullingham CI, Miller JM, Peery RM, Dupuis JR, Malenfant RM, Gorrell JC, Janes JK. 2020. Confidently identifying the correct K value using the ΔK method: When does K = 2? Molecular Ecology 29(5): 862–869.

Dane F, Lang P, Huang H, Fu Y. 2003. Intercontinental genetic divergence of *Castanea* species in eastern Asia and eastern North America. Heredity 91(3): 314–321.

Dering M, Baranowska M, Beridze B, Chybicki IJ, Danelia I, Iszkuło G, Kvartskhava G, Kosiński P, Rączka G, Thomas PA, Tomaszewski D. 2021. The evolutionary heritage and ecological uniqueness of Scots pine in the Caucasus ecoregion is at risk of climate changes. Scientific Reports 11(1): 1–17.

Dumbadze G, Termel G, Vasadze T, Lomtatidze N, Chakhvadze K. 2018. Intensity of Chestnut drying and natural restoration of forest in Keda municipality (Ajara, Georgia). International Journal of Ecosystem and Ecology Science 8: 347–352.

Dumolin S, Demesure B, Petit RJ. 1995. Inheritance of chloroplast and mitochondrial genomes in pedunculate oak investigated with an efficient PCR method. Theoretical and Applied Genetics 91: 1253–1256.

Ekhavaia J, Simeone MC, Silakadze N, Abdaladze O. 2018. Morphological diversity and phylogeography of the Georgian durmast oak (*Q. petraea* subsp. iberica) and related Caucasian oak species in Georgia (South Caucasus). TGG 14(2): 17.

Evanno G, Regnaut S, Goudet J. 2005. Detecting the number of clusters of individuals using the software STRUCTURE: a simulation study. Molecular Ecology 14(8): 2611–2620.

Fineschi S, Taurchini D, Villani F, Vendramin GG. 2000. Chloroplast DNA polymorphism reveals little geographical structure in *Castanea sativa* Mill. (Fagaceae) throughout southern European countries. Molecular Ecology 9: 1495–1503.

Fordham DA, Saltré F, Haythorne S, Wigley TM, Otto-Bliesner BL, Chan KC, Brook B.W. 2017. PaleoView: a tool for generating continuous climate projections spanning the last 21 000 years at regional and global scales. Ecography 40(11): 1348–1358.

Freitas TR, Santos JA, Silva AP, Fraga H. 2021. Influence of climate change on chestnut trees: A review. Plants 10(7): 1463.

Grímsson F, Zetter R, Grimm GW, Pedersen GK, Pedersen AK, Denk T. 2015. Fagaceae pollen from the early Cenozoic of West Greenland: revisiting Engler’s and Chaney’s Arcto-Tertiary hypotheses. Plant Systematics and Evolution 301(2): 809–832.

Goudet J. 1995. FSTAT (version 1.2): a computer program to calculate F-statistics. Journal of heredity 86(6): 485–486.

Head MJ, Gibbard PL. 2015. Early–Middle Pleistocene transitions: Linking terrestrial and marine realms. Quaternary International 389: 7–46.

IPCC, 2014: Climate Change 2014: Synthesis Report. Contribution of Working Groups I, II and III to the Fifth Assessment Report of the Intergovernmental Panel on Climate Change [Core Writing Team, R.K. Pachauri and L.A. Meyer (eds.)]. IPCC, Geneva, Switzerland, 151 pp. Figure SPM.7

IPCC, 2021: Climate Change 2021: The Physical Science Basis. Contribution of Working Group I to the Sixth Assessment Report of the Intergovernmental Panel on Climate Change [Masson-Delmotte, V., P. Zhai, A. Pirani, S.L. Connors, C. Péan, S. Berger, N. Caud, Y. Chen, L. Goldfarb, M.I. Gomis, M. Huang, K. Leitzell, E. Lonnoy, J.B.R. Matthews, T.K. Maycock, T. Waterfield, O. Yelekçi, R. Yu, and B. Zhou (eds.)]. Cambridge University Press, Cambridge, United Kingdom and New York, NY, USA, In press, doi:10.1017/9781009157896.

Janfaza S, Nasr SMH, Yousefzadeh H, Botta R. 2015. Phylogenetic relationships of the genus Castanea based on chloroplast rbcl with focusing Iranian chestnut. Journal of Biodiversity and Ecological Sciences 5: 312–323.

Janfaza S, Yousefzadeh H, Nasr SMH, Botta R, Abkenar AA, Marinoni DT. 2017. Genetic diversity of *Castanea sativa* an endangered species in the Hyrcanian forest. Silva Fennica 51: 1–15.

Jia D-R, Abbott RJ, Liu T-L, Mao K-S, Bartisch IV, Liu J-Q. 2012. Out of the Qinghai–Tibet Plateau: evidence for the origin and dispersal of Eurasian temperate plants from a phylogeographic study of *Hippophaë rhamnoides* (Elaeagnaceae). New Phytologist 194: 1123–1133.

Karger DN, Conrad O, Böhner J, Kawohl T, Kreft H, Soria-Auza RW, Zimmermann NE, Linder P, Kessler M. 2017. Climatologies at high resolution for the Earth land surface areas. Scientific Data 4(1): 1–20.

Kadereit JW, Licht W, Uhnik ChH. 2010. Asian relationships of the flora of the European Alps. Plant Ecology and Diversity 1:2: 171–179.

Koch MA, Michling F, Walther A, Huang XC, Tewes L, Müller C. 2017. Early-Mid Pleistocene genetic differentiation and range expansions as exemplified by invasive Eurasian *Bunias orientalis* (Brassicaceae) indicates the Caucasus as key region. Scientific Reports 7: 16764.

Krijgsman W, Tesakov A, Yanina T, Lazarev S, Danukalova G, Van Baak CGC, Agustí J, Alçiçek MC, Aliyeva E, Bista D, Bruch A, Büyükmeriç Y, Bukhsianidze M, Flecker R, Frolov P, Hoyle TM, Jorissen EL, Kirscher U, Koriche SA, Kroonenberg SB, Lordkipanidze D, Oms O, Rausch L, Singarayer J, Stoica M, van de Velde S, Titov VV, Wesselingh FP. 2019. Quaternary time scales for the Pontocaspian domain: Interbasinal connectivity and faunal evolution. Earth-Science Reviews 188: 1–40.

Kalinowski ST, Taper ML. 2006. Maximum likelihood estimation of the frequency of null alleles at microsatellite loci. Conservation Genetics 7(6): 991–995.

Krebs P, Conedera M, Pradella M, Torriani D, Felber M, Tinner W. 2004. Quaternary refugia of the sweet chestnut (*Castanea sativa* Mill.): an extended palynological approach. Vegetation history and Archaeobotany 13(3): 145–160.

Krebs P, Pezzatti GB, Beffa G, Tinner W, Conedera, M. 2019. Revising the sweet chestnut (*Castanea sativa* Mill.) refugia history of the last glacial period with extended pollen and macrofossil evidence. Quaternary Science Reviews 206: 111–128.

Kvaček Z, & Walther H. 2001. The Oligocene of Central Europe and the development of forest vegetation in space and time based on megafossils. Palaeontographica Abteilung B, 259, 125–148.

Lang P, Dane F, Kubisiak TL. 2006. Phylogeny of *Castanea* (Fagaceae) based on chloroplast trnT-LF sequence data. Tree Genetics & Genomes 2(3): 132–139.

Larue C, Austruy E, Basset G, Petit RJ. 2021. Revisiting pollination mode in chestnut (*Castanea* spp.): an integrated approach. Botany Letters 168: 348–372.

Leroy SA, Arpe K. 2007. Glacial refugia for summer-green trees in Europe and south-west Asia as proposed by ECHAM3 time-slice atmospheric model simulations. Journal of Biogeography 34(12): 2115–2128.

Levin BA, Gandlin AA, Simonov ES, Levina MA, Barmintseva AE, Japoshvili B, Mugue NS, Mumladze L, Mustafayev NJ, Pashkov AN, Roubenyan HR, Shapovalov MI, Doadrio I. 2019. Phylogeny, phylogeography and hybridization of Caucasian barbels of the genus *Barbus* (Actinopterygii, Cyprinidae). Molecular Phylogenetics and Evolution 135: 31–44.

Li YL, Liu JX. 2018. StructureSelector: A web based software to select and visualize the optimal number of clusters using multiple methods. Molecular Ecology Resources 18:176–177

Lisiecki LE, Raymo ME. 2005. A Pliocene-Pleistocene stack of 57 globally distributed benthic δ^18^ O records. Paleoceanography 20: 1–17.

Maharramova EH, Safarov HM, Kozlowski G, Borsch T, Muller LA. 2015. Analysis of nuclear microsatellites reveals limited differentiation between Colchic and Hyrcanian populations of the wind-pollinated relict tree *Zelkova carpinifolia* (Ulmaceae). American Journal of Botany 102(1): 119–128.

Magri D, Polombo MR. 2013. Early to Middle Pleistocene dynamics of plant and mammal communities in South West Europe. Quaternary international: the journal of the International Union for Quaternary Research 288: 63–72.

Mahler S, Shatilova I, Bruch A. 2022. Neogene long-term trends in climate of the Colchic vegetation refuge in Western Georgia – Uplift versus global cooling. Review of Palaeobotany and Palynology 296: 104546.

Manafzadeh S, Staedler YM, Conti E. 2017. Visions of the past and dreams of the future in the Orient: the Irano-Turanian region from classical botany to evolutionary studies. Biological Reviews of the Cambridge Philosophical Society 92: 1365–1388.

Marinoni D, Akkak A, Bounous G, Edwards KJ, Botta R. 2003. Development and characterization of microsatellite markers in *Castanea sativa* (Mill.). Molecular Breeding 11(2): 127–136.

Martín MA, Angela Martín M, Mattioni C, Molina JR, Alvarez JB, Cherubini M, Herrera MA, Villani F, Martín LM. 2012. Landscape genetic structure of chestnut (*Castanea sativa* Mill.) in Spain. Tree Genetics & Genomes 8: 127–136.

Mattioni C, Martin MA, Pollegioni P, Cherubini M, Villani F. 2013. Microsatellite markers reveal a strong geographical structure in European populations of *Castanea sativa* (Fagaceae): evidence for multiple glacial refugia. American Journal of Botany 100: 951–961.

Mattioni C, Martin MA, Chiocchini F, Cherubini M, Gaudet M, Pollegioni P, Velichkov I, Jarman R, Chambers FM, Paule L, Damian VL. 2017. Landscape genetics structure of European sweet chestnut (*Castanea sativa* Mill): indications for conservation priorities. Tree Genetics & Genomes 13(2): 1–14.

Mayol M, Riba M, González-Martínez SC, Bagnoli F, de Beaulieu JL, Berganzo E, Burgarella C, Dubreuil M, Krajmerová D, Paule L, Romšáková I, Vettori C, Vincenot L, Vendramin GG. 2015. Adapting through glacial cycles: insights from a long-lived tree (*Taxus baccata*). The New Phytologist 208: 973–986.

Mercuri AM, Bandini Mazzanti M, Florenzano A, Montecchi MC, Rattighieri E. 2013. Olea, Juglans and Castanea: The OJC group as pollen evidence of the development of human-induced environments in the Italian peninsula. Quaternary International 303: 24–42.

Meirmans PG. 2015. Seven common mistakes in population genetics and how to avoid them. Molecular Ecology 24(13): 3223–3231.

Messager E, Poulenard J, Sabatier P, Develle AL, Wilhelm B, Nomade S, Scao V, Giguet-Covex C, Von Grafenstein U, Arnaud F, Malet E, Mgeladze A, Herrscher E, Banjan M, Mazuy A, Dumoulin JP, Belmecheri S, Lordkipanidze D. 2021. Paravani, a puzzling lake in the South Caucasus. Quaternary International 579: 6–18.

Milne RI, Abbott RJ. 2002. The origin and evolution of Tertiary relict floras. Advances in Botanical Research 38: 281–317.

Mittermeier RA, Turner WR, Larsen FW, Brooks TM, Gascon C. 2011. Global biodiversity conservation: the critical role of hotspots. In: Zachos FE, Habel JC, eds. Biodiversity hotspots. Berlin, Heidelberg: Springer, 3–22.

Mosbrugger V, Utescher T, Dilcher DL. 2005. Cenozoic continental climatic evolution of Central Europe. PNAS 102: 14964–14969.

Myers N, Mittermeier, RA, Mittermeier CG, Da Fonseca GA, Kent J. 2000. Biodiversity hotspots for conservation priorities. Nature 403(6772): 853–858.

Naimi B, Hamm NA, Groen TA, Skidmore AK, Toxopeus AG. 2014. Where is positional uncertainty a problem for species distribution modelling? Ecography 37(2): 191–203.

Nakhutsrishvili G, Zazanashvili N, Batsatsashvili K, Montalvo Mancheno CS. 2015. Colchic and Hyrcanian forests of the Caucasus: similarities, differences and conservation status. Flora Mediterranea 25: 185–192.

Nakhutsrishvili G, Abdaladze O. 2017. Plant diversity of the central Great Gaugasus. In: Plant diversity in the Central Great Caucasus: a quantitative assessment. Springer, 17–111.

Neiber MT, Hausdorf B. 2015. Phylogeography of the land snail genus *Circassina* (Gastropoda: Hygromiidae) implies multiple Pleistocene refugia in the western Caucasus region. Molecular Phylogenetics and Evolution 93: 129–142.

Olson DM, Dinerstein E, Wikramanayake ED, Burgess ND, Powell GV, Underwood EC, D’amico JA, Itoua I, Strand HE, Morrison JC, Loucks CJ. 2001. Terrestrial Ecoregions of the World: A New Map of Life on Earth: A new global map of terrestrial ecoregions provides an innovative tool for conserving biodiversity. BioScience 51(11): 933–938.

Orain R, Russo Ermolli E, Lebreton V, Di Donato V, Bahain JJ, Sémah AM. 2015. Vegetation sensitivity to local environmental factors and global climate changes during the Middle Pleistocene in southern Italy—A case study from the Molise Apennines. Review of Palaeobotany and Palynology 220: 69–77.

Orsini L, Vanoverbeke J, Swillen I, Mergeay J, De Meester L. 2013. Drivers of population genetic differentiation in the wild: isolation by dispersal limitation, isolation by adaptation and isolation by colonization. Molecular Ecology 22: 5983–5999.

Otto-Bliesner BL, Marshall SJ, Overpeck JT, Miller GH, Hu A, CAPE Last Interglacial Project members. 2006. Simulating Arctic climate warmth and icefield retreat in the last interglaciation. science 311(5768): 1751–1753.

Parvizi E, Keikhosravi A, Naderloo R, Solhjouy-Fard S, Sheibak F, Schubart CD. 2019. Phylogeography of Potamon ibericum (Brachyura: Potamidae) identifies Quaternary glacial refugia within the Caucasus biodiversity hot spot. Ecology and Evolution 9(8): 4749–4759.

Peper J, Pietzsch D, Manthey M. 2010. Semi-arid rangeland vegetation of the Greater Caucasus foothills in Azerbaijan and its driving environmental conditions. Phytocoenologia 40(2-3): 73–90.

Peakall R, Smouse PE. 2012. GenAlEx 6.5: genetic analysis in Excel. Population genetic software for teaching and research-an update. Bioinformatics 28: 2537–2539.

Phillips SJ, Dudík M, Schapire RE. 2004. A maximum entropy approach to species distribution modeling. In Proceedings of the Twenty-First International Conference on Machine Learning 83 (Association for Computing Machinery).

Pokryszko BM, Cameron RA, Mumladze L, Tarkhnishvili D. 2011. Forest snail faunas from Georgian Transcaucasia: patterns of diversity in a Pleistocene refugium. Biological Journal of the Linnean Society 102(2): 239–250.

Pollegioni P, Lungo SD, Müller R, Woeste KE, Chiocchini F, Clark J, Hemery GE, Mapelli S, Villani F, Malvolti ME, Mattioni C. 2020. Biocultural diversity of common walnut (*Juglans regia* L.) and sweet chestnut (*Castanea sativa* Mill.) across Eurasia. Ecology and Evolution 10(20): 11192–11216.

Popescu SM, Biltekin D, Winter H, Suc JP, Melinte-Dobrinescu MC, Klotz S, Rabineau M, Combourieu-Nebout N, Clauzon G, Deaconu F. 2010. Pliocene and Lower Pleistocene vegetation and climate changes at the European scale: Long pollen records and climatostratigraphy. Quaternary International 219(1-2): 152–167.

Popov SV, Shcherba IG, Ilyina L B, Nevesskaya LA. Paramonova N P, Khondkarin OS, Magyar I. 2006. Late Miocene to Pliocene paleogeography of the Paratethys and its relation to the Mediterranean. Palaeogeography, Palaeoclimatology, Palaeoecology 238: 91–106.

Postigo-Mijarra JM, Morla C, Barrón E, Morales-Molino C, García S. 2010. Patterns of extinction and persistence of Arctotertiary flora in Iberia during the Quaternary. Review of Palaeobotany and Palynology 162: 416–426.

Pridnya MV, Cherpakov VV, Paillet FL. 1996. Ecology and Pathology of European Chestnut (*Castanea sativa*) in the Deciduous Forests of the Caucasus Mountains in Southern Russia. Bulletin of the Torrey Botanical Club 123: 213.

Pritchard JK, Stephens M, Donnelly P. 2000. Inference of population structure using multilocus genotype data. Genetics 155: 945–959.

Prospero S, Lutz A, Tavadze B, Supatashvili A, Rigling D. 2013. Discovery of a new gene pool and a high genetic diversity of the chestnut blight fungus *Cryphonectria parasitica* in Caucasian Georgia. Infection, Genetics and Evolution 20: 131–139.

Pudlo P, Marin JM, Estoup A, Cornuet JM, Gautier M, Robert CP. 2016. Reliable ABC model choice via random forests. Bioinformatics 32: 859–866.

Puechmaille SJ. 2016. The program structure does not reliably recover the correct population structure when sampling is uneven: subsampling and new estimators alleviate the problem. Molecular Ecology Resources 16(3): 608–627.

QGIS.org, 2021. QGIS Geographic Information System. QGIS Association. http://www.qgis.org

Raymond M, Rousset F. 1995. GENEPOP (version 1.2): population genetics software for exact tests and ecumenicism. Journal of Heredity 86: 248–249

Roces-Díaz JV, Jiménez-Alfaro B, Chytrý M, Díaz-Varela ER, Álvarez-Álvarez P. 2018. Glacial refugia and mid-Holocene expansion delineate the current distribution of *Castanea sativa* in Europe. Palaeogeography, Palaeoclimatology, Palaeoecology 491: 152–160.

Rousset F. 2008. Genepop’007: a complete reimplementation of the Genepop software for Windows and Linux. Molecular Ecology Resources 8: 103–106

Ryabogina NE, Afonin AS, Ivanov SN, Li HC, Kalinin PA, Udaltsov SN, Nikolaenko SA. 2019. Holocene paleoenvironmental changes reflected in peat and lake sediment records of Western Siberia: Geochemical and plant macrofossil proxies. Quaternary International 528: 73–87.

Sękiewicz K, Danelia I, Farzaliyev V, Gholizadeh H, Iszkuło G, Naqinezhad A, Ramezani E, Thomas PA, Tomaszewski D, Walas Ł, Dering M. 2022. Past climatic refugia and landscape resistance explain spatial genetic structure in Oriental beech in the South Caucasus. Ecology and Evloution 12: e9320.

Scotti-Saintagne C, Boivin T, Suez M, Musch B, Scotti I, Fady B. 2021. Signature of mid-Pleistocene lineages in the European silver fir (*Abies alba* Mill.) at its geographic distribution margin. Ecology and Evolution 11(16): 10984–10999.

Shackleton S, Menking JA, Brook E, Buizert Ch, Dyonisius MN, Petrenko VV, Baggenstos D, Severinghaus JP. 2021. Evolution of mean ocean temperature in Marine Isotope Stage 4. Climate of the Past 17: 2273–2289.

Shatilova I, Mchedlishvili N, Rukhadze L, Kvavadze E. 2011. The history of the flora and vegetation of Georgia. Lordkipanidze D, Vekua A, eds. Tbilisi: Georgian national museum, Institute of paleobiology.

Shatilova I, Kokolashvili IM, Bukhsianidze M.G, Koiava KP, Maissuradze LS, Bruch AA. 2021. Late Cenozoic bioevents on the territory of Georgia (foraminifera and pollen). Georgian National Museum, Tbilisi, Georgia. 160 pp.

Song YG, Walas Ł, Pietras M, Sâm HV, Yousefzadeh H, Ok T, Farzaliyev V, Worobiec G, Worobiec E, Stachowicz-Rybka R, Boratyński A. 2021. Past, present and future suitable areas for the relict tree *Pterocarya fraxinifolia* (Juglandaceae): Integrating fossil records, niche modeling, and phylogeography for conservation. European Journal of Forest Research 140(6): 1323–1339.

Sosson M, Rolland Y, Müller C, Danelian T, Melkonyan R, Kekelia S, Adamia S, Babazadeh V, Kangarli T, Avagyan A, Galoyan G. 2010. Subductions, obduction and collision in the Lesser Caucasus (Armenia, Azerbaijan, Georgia), new insights. Geological Society, London, Special Publications 340(1): 329–352.

Tarkhnishvili D, 2014. Historical biogeography of the Caucasus. NOVA Science Publishers.

Tarkhnishvili D, Gavashelishvili A, Mumladze L. 2012. Palaeoclimatic models help to understand current distribution of Caucasian forest species. Biological Journal of the Linnean Society 105: 231–248.

Tavadze B L, Supatashvili AS, Rigling D, Sotirovski K, Mamukashvili TI, Chitia ST. 2013. Pathological status of chestnut stands in Georgia. Annals of Agrarian Science 11(1): 104–107.

Thomas PA, Alhamed O, Iszkuło G, Dering M, Mukassabi T. 2019. Biological Flora of the British Isles: *Aesculus hippocastanum*. Journal of Ecology 107(2): 992–1030.

Tzedakis PC, Hooghiemstra H, Pälike H. 2006. The last 1.35 million years at Tenaghi Philippon: revised chronostratigraphy and long-term vegetation trends. Quaternary Science Reviews 25(23-24): 3416–3430.

Villani F, Pigliucci M, Cherubini M. 1994. Evolution of *Castanea sativa* Mill. in Turkey and Europe. Genet Resources 63: 109–116.

Volkova P, Laczkó L, Demina O, Schanzer I, Sramkó G. 2020. Out of Colchis: the colonization of Europe by *Primula vulgaris* Huds. (Primulaceae). Acta Societatis Botanicorum Poloniae 89(3): Article-89313.

Walas Ł, Ganatsas P, Iszkuło G, Thomas PA, Dering M. 2019. Spatial genetic structure and diversity of natural populations of *Aesculus hippocastanum* L. in Greece. PlosOne 14(12): e0226225.

Wang Y, Xie B, Wan F, Xiao Q. 2007. Application of ROC curve analysis in evaluating the performance of alien species’ potential distribution models. Biodiversity Science 15(4): 365–372.

Yazvenko SB. 1994. Holocene pollen record from a peatland in the West Caucasus, Abkhasia, Black Sea region. Journal of Paleolimnology 12: 65–74.

Yousefzadeh H, Hosseinzadeh Colagar A, Akbarzadeh F, Tippery NP. 2014. Taxonomic status and genetic differentiation of Hyrcanian Castanea based on noncoding chloroplast DNA sequences data. Tree Genetics & Genomes 10(6): 1611–1629.

Zazanashvili N, Gavashelishvili L, Montalvo C, Beruchashvili G, Heidelberg A, Neuner J, Schulzke R, Garforth M. 2011. Strategic Guidelines for Responding to Impacts of Global Climate Change on Forests in the Southern Caucasus (Armenia, Azerbaijan, Georgia). WWF, KfW.

Zazanashvili N, Sanadiradze G, Bukhnikashvili A, Kandaurov A, Tarkhnishvili D. 2004. Caucasus. In: Mittermaier RA, Gil PG, Hoffmann M, Pilgrim J, Brooks T, Mittermaier CG, Lamoreux J, da Fonseca GAB, eds. Hotspots revisited, Earth’s biologically richest and most endangered terrestrial ecoregions. Sierra Madre: CEMEX/Agrupacion Sierra Madre, 148–153.

Zohary M. 1973. Geobotanical Foundations of the Middle East (Volume I and II). Gustav Fischer Verlag, Stuttgart and Swets & Zeitlinger, Amsterdam.

Zohary D, Hopf M, Weiss E. 2012 Domestication of Plants in the Old World: The origin and spread of domesticated plants in Southwest Asia, Europe, and the Mediterranean Basin. Oxford University Press.

Zyryanov EV. 1992. Palynological investigations of Miocene deposits on the New Siberian Archipelago (USSR). Arctic 45(3): 285–294.

