## Supplementary Material 1 for "Evolutionary history of *Castanea sativa* Mill. in the Caucasus driven by Middle and Late Pleistocene paleoenvironmental changes"

Suplementary Material 1

**Demographic investigations performed with ABC-RF approach**

ORCID:

BB: 0000-0003-4189-5392

KS: 0000-0002-1830-0816

ŁW: 0000-0002-4060-9801

PT: 0000-0003-3115-3301

GK: 0000-0002-5654-4061

VF: 0000-0002-4004-234X

AB: 0000-0002-4629-1507

MD: 0000-0001-7017-5541

Table S1. Priors used for ABC procedure in DIYABC Random Forest for demographic history inferences

| **Parameters** | **Distribution** | **Min-Max** | **Mean** | **Shape** |
| --- | --- | --- | --- | --- |
| **Genetic parameters** |  |  |  |  |
| Mean mutation rate | Log uniform | 1.10^-5^-1.10^-3^ |  |  |
| Individual mutation rate | Gamma | 1.10^-7^-1.10^-1^ | Mean mutation rate | 2 |
| Mean coefficient P | Uniform | 0.1-12 |  |  |
| Individual locus coefficient P | Gamma | 0.1-15 | Mean coefficient P | 2 |
| **Historical parameters** | | | | |
| Lineage 1 |  | 10-10000 |  |  |
| Lineage 2 |  | 10-1000 |  |  |
| Lineage 3 |  | 10-10000 |  |  |
| Lineage 4 |  | 10-7000 |  |  |
| Lineage 5 |  | 10-10000 |  |  |
| *ta* |  | 10-2000 |  |  |
| *ra* |  | 0.001-0.999 |  |  |
| *t1* |  | 10-5000 |  |  |
| *t2* |  | 10-3000 |  |  |
| *t3* |  | 10-10000 |  |  |
| *t4* |  | 10-10000 |  |  |
| Conditions | t1>ta, t2>ta, t2>t1, t3>t2, t4>t3 | | | |

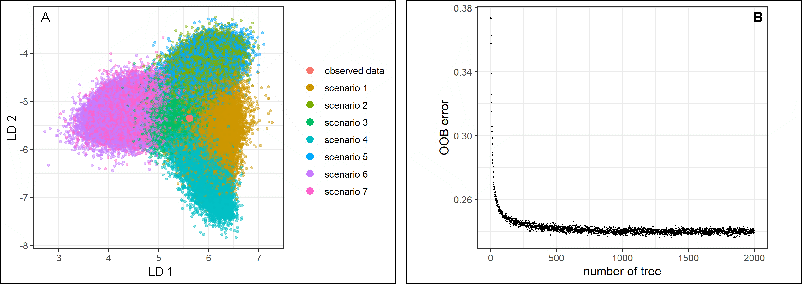

Fig. S1. Scenario choice based on the Random Forest approach implemented in DIYABC Random Forest: (A) Linear discriminant analysis (LDA) projection of the datasets from the training set and observed data on the two first LD axis; (B) prediction power indicating global prior error rate computed using out of bootstrap (OOB error, set as 1000) for random forest versus the number of growing trees in the forest.

Table S2. Posterior probability (PP) obtained for seven tested demographic scenarios of divergence tested with Random Forest approach implemented in DIYABC-Random Forest. Scenario 1 was indicated as having the highest posterior probability and so was assumed as the most probable one for populations of *C. sativa* in the Caucasus. Scenario choice was performed in ten replicates ABC-RF analyses based on 140,040 simulated training datasets. The table also presents the accuracy metrics of prediction. The number of trees in the constructed random forests was set to 2,000. Standard deviations over the 10 replicate analyses are given in brackets for each metrics. Scenarios are depicted in Figure 2 in the main text of the article.

| **ABC-FR run** | **Scenario 1** | **Scenario 2** | **Scenario 3** | **Scenario 4** | **Scenario 5** | **Scenario 6** | **Scenario 7** | **Scenario choice** | **Global error**  **rate** | **Local error**  **rate** | **Posterior probability** |
| --- | --- | --- | --- | --- | --- | --- | --- | --- | --- | --- | --- |
| 1 | 616 | 220 | 318 | 226 | 204 | 212 | 204 | 1 | 0.242095 | 0.290 | 0.710 |
| 2 | 644 | 226 | 266 | 257 | 189 | 220 | 198 | 1 | 0.242252 | 0.290 | 0.710 |
| 3 | 631 | 243 | 288 | 240 | 190 | 227 | 181 | 1 | 0.241924 | 0.285 | 0.715 |
| 4 | 631 | 236 | 272 | 260 | 172 | 232 | 197 | 1 | 0.241681 | 0.280 | 0.720 |
| 5 | 679 | 229 | 246 | 232 | 190 | 220 | 204 | 1 | 0.242252 | 0.271 | 0.729 |
| 6 | 632 | 224 | 280 | 226 | 217 | 231 | 190 | 1 | 0.242416 | 0.269 | 0.731 |
| 7 | 634 | 232 | 278 | 247 | 194 | 212 | 203 | 1 | 0.242416 | 0.277 | 0.723 |
| 8 | 654 | 236 | 248 | 242 | 194 | 229 | 197 | 1 | 0.242379 | 0.263 | 0.737 |
| 9 | 667 | 238 | 245 | 233 | 188 | 222 | 207 | 1 | 0.242437 | 0.267 | 0.733 |
| 10 | 675 | 224 | 251 | 240 | 206 | 200 | 204 | 1 | 0.242495 | 0.284 | 0.716 |
| ***Average*** | 646  *(21.417)* | 231  *(7.391)* | 269  *(23.218)* | 240  *(11.767)* | 194  *(12.240)* | 221  *(10.135)* | 199  *(7.934)* | **1** | 0.242  *(0.0003)* | 0.278  *(0.0097)* | 0.722  *(0.010)* |
