## Supplementary Material 2 for "Evolutionary history of *Castanea sativa* Mill. in the Caucasus driven by Middle and Late Pleistocene paleoenvironmental changes"

Suplementary Material 2

**MaxEnt analysis on species distribution modelling**

ORCID:

BB: 0000-0003-4189-5392

KS: 0000-0002-1830-0816

ŁW: 0000-0002-4060-9801

PT: 0000-0003-3115-3301

GK: 0000-0002-5654-4061

VF: 0000-0002-4004-234X

AB: 0000-0002-4629-1507

MD: 0000-0001-7017-5541

**Table S1.** Coordinates of sites of *C. sativa* used for SDMs of species in the South Caucasus.

|  | Latitude | Longitude | Source |
| --- | --- | --- | --- |
| 1 | 41.9326 | 46.0194 | Studied populations in this work |
| 2 | 42.3557 | 42.9991 | Studied populations in this work |
| 3 | 42.3329 | 42.9421 | Studied populations in this work |
| 4 | 42.5038 | 43.1462 | Studied populations in this work |
| 5 | 42.5205 | 43.1896 | Studied populations in this work |
| 6 | 42.3442 | 43.5264 | Studied populations in this work |
| 7 | 41.8853 | 46.2457 | Studied populations in this work |
| 8 | 42.0437 | 43.4988 | Studied populations in this work |
| 9 | 42.1429 | 43.1068 | Studied populations in this work |
| 10 | 42.6548 | 42.2216 | Studied populations in this work |
| 11 | 41.9581 | 42.7697 | Studied populations in this work |
| 12 | 42.6569 | 42.4341 | Studied populations in this work |
| 13 | 41.9293 | 42.3735 | Studied populations in this work |
| 14 | 41.7287 | 42.0781 | Studied populations in this work |
| 15 | 41.6843 | 41.8345 | Studied populations in this work |
| 16 | 42.2226 | 45.3037 | Studied populations in this work |
| 17 | 41.6796 | 46.6864 | Studied populations in this work |
| 18 | 41.6194 | 46.6990 | Studied populations in this work |
| 19 | 40.9883 | 47.9544 | Studied populations in this work |
| 20 | 41.2976 | 47.1249 | Studied populations in this work |
| 21 | 41.5561 | 41.9899 | National Botanical Garden of Georgia |
| 22 | 41.9067 | 42.7408 | National Botanical Garden of Georgia |
| 23 | 42.7754 | 42.0791 | National Botanical Garden of Georgia |
| 24 | 41.8935 | 42.2980 | National Botanical Garden of Georgia |
| 25 | 41.0273 | 40.6062 | gbif, https://doi.org/10.15468/dl.5nqse6 |
| 26 | 41.5704 | 41.8461 | gbif, https://doi.org/10.15468/dl.5nqse6 |
| 27 | 42.0973 | 43.4009 | gbif, https://doi.org/10.15468/dl.5nqse6 |
| 28 | 43.0340 | 41.4686 | gbif, https://doi.org/10.15468/dl.5nqse6 |
| 29 | 43.0936 | 41.6781 | gbif, https://doi.org/10.15468/dl.5nqse6 |
| 30 | 43.4201 | 40.1146 | gbif, https://doi.org/10.15468/dl.5nqse6 |
| 31 | 43.9158 | 40.1420 | gbif, https://doi.org/10.15468/dl.5nqse6 |
| 32 | 43.9998 | 40.1543 | gbif, https://doi.org/10.15468/dl.5nqse6 |
| 33 | 44.0500 | 40.1667 | gbif, https://doi.org/10.15468/dl.5nqse6 |
| 34 | 44.2638 | 39.2739 | gbif, https://doi.org/10.15468/dl.5nqse6 |
| 35 | 44.3506 | 39.1892 | gbif, https://doi.org/10.15468/dl.5nqse6 |
| 36 | 43.6737 | 40.2983 | iNaturalist, <https://www.inaturalist.org/observations/33945669> |
| 37 | 41.2766 | 41.3754 | iNaturalist, <https://www.inaturalist.org/observations/38457989> |
| 38 | 41.0780 | 47.4611 | iNaturalist, <https://www.inaturalist.org/observations/7739469> |
| 39 | 42.6414 | 43.5333 | iNaturalist, <https://www.inaturalist.org/observations/31516200> |
| 40 | 44.1671 | 39.0524 | iNaturalist, <https://www.inaturalist.org/observations/57429598> |
| 41 | 43.9022 | 39.8360 | iNaturalist, <https://www.inaturalist.org/observations/48557093> |
| 42 | 43.7075 | 39.7789 | iNaturalist, <https://www.inaturalist.org/observations/57446646> |
| 43 | 43.6904 | 40.2294 | iNaturalist, <https://www.inaturalist.org/observations/57890566> |
| 44 | 43.5308 | 39.8778 | iNaturalist, <https://www.inaturalist.org/observations/44954472> |
| 45 | 43.0911 | 40.8207 | iNaturalist, <https://www.inaturalist.org/observations/41242964> |
| 46 | 44.3350 | 39.0514 | iNaturalist, <https://www.inaturalist.org/observations/78566210> |
| 47 | 44.2469 | 39.2566 | iNaturalist, <https://www.inaturalist.org/observations/62028748> |
| 48 | 44.0183 | 39.3730 | iNaturalist, <https://www.inaturalist.org/observations/75068721> |
| 49 | 43.6190 | 40.0447 | iNaturalist, <https://www.inaturalist.org/observations/66563138> |
| 50 | 43.6528 | 40.0662 | iNaturalist, <https://www.inaturalist.org/observations/66774984> |
| 51 | 43.6450 | 40.3139 | iNaturalist, <https://www.inaturalist.org/observations/60871603> |
| 52 | 41.8423 | 46.2827 | iNaturalist, <https://www.inaturalist.org/observations/80074379> |
| 53 | 42.0963 | 43.1626 | iNaturalist, <https://www.inaturalist.org/observations/15697496> |
| 54 | 42.3988 | 43.3629 | iNaturalist, <https://www.inaturalist.org/observations/80874547> |
| 55 | 41.4661 | 47.0548 | iNaturalist, <https://www.inaturalist.org/observations/66016276> |
| 56 | 43.7334 | 40.0349 | Mattioni *et al.,* 2017 |
| 57 | 42.7439 | 41.4363 | Mattioni *et al.,* 2017 |
| 58 | 42.8086 | 41.7747 | Mattioni *et al., 2017* |
| 59 | 42.0524 | 43.4926 | Mattioni *et al.,* 2017 |
| 60 | 41.6833 | 41.8667 | Mattioni *et al.,* 2017 |
| 61 | 40.7200 | 39.6300 | Mattioni *et al.,* 2017 |
| 62 | 41.5709 | 41.8587 | Private observation |
| 63 | 41.9832 | 43.3613 | Private observation |
| 64 | 41.9863 | 43.3489 | Private observation |
| 65 | 42.2503 | 43.2278 | Private observation |
| 66 | 42.2422 | 43.2330 | Private observation |
| 67 | 43.2577 | 41.0858 | Private observation |
| 68 | 41.0759 | 47.7299 | Private observation |
| 69 | 41.0958 | 47.5543 | Private observation |
| 70 | 41.2212 | 47.2978 | Private observation |
| 71 | 41.6105 | 46.7610 | Private observation |
| 72 | 41.0554 | 47.7989 | Private observation |
| 73 | 40.9822 | 47.9078 | Private observation |
| 74 | 41.9763 | 42.4074 | Private observation |
| 75 | 41.7572 | 41.9788 | Private observation |
| 76 | 41.8541 | 46.2873 | Private observation |
| 77 | 41.4873 | 41.8020 | Private observation |
| 78 | 41.4957 | 41.8759 | Private observation |
| 79 | 41.4893 | 41.7470 | Private observation |
| 80 | 42.5234 | 42.4489 | Private observation |
| 81 | 42.5238 | 42.4511 | Private observation |
| 82 | 41.0258 | 41.0120 | Private observation |
| 83 | 40.6824 | 39.6590 | Private observation |
| 84 | 40.8000 | 39.6716 | Private observation |
| 85 | 40.9242 | 39.7383 | Private observation |
| 86 | 40.8367 | 40.4804 | Private observation |
| 87 | 41.0812 | 41.0356 | Private observation |
| 88 | 41.3817 | 41.5692 | Private observation |
| 89 | 37.2667 | 49.2500 | Janfaza *et al.,* 2017 |
| 90 | 37.5000 | 49.0333 | Janfaza *et al.,* 2017 |
| 91 | 37.0833 | 49.2333 | Janfaza *et al.,* 2017 |
| 92 | 37.1833 | 49.4667 | Janfaza *et al.,* 2017 |

Private observations category includes the record gained from local foresters.

Mattioni C, Martin MA, Chiocchini F, Cherubin M, Gaudet M, Pollegioni , Velichkov I, Jarman R, Chambers FM, Paule L & Damian VL. 2017. Landscape genetics structure of European sweet chestnut (Castanea sativa Mill): indications for conservation priorities. *Tree Genetics & Genomes* 13: 1-14.

Janfaza S, Yousefzadeh H, Hosseini Nasr, SM, Botta R, Asadi Abkenar A & Torello Marinoni D. 2017. Genetic diversity of *Castanea sativa* an endangered species in the Hyrcanian forest. *Silva Fennica* 51:e1705

**Table S2.** The contribution of eight non-corralated bioclimatic varaibles.

| Code | Bioclimatic variable | LIG | LGM | EH | MH | LH | Current |
| --- | --- | --- | --- | --- | --- | --- | --- |
| Bio01 | Annual Mean Temperature | **13.2** | **12.5** | **12.5** | **12.4** | **11.6** | **11.6** |
| Bio03 | Isothermality | 0.2 | 0.4 | 0.6 | 0.4 | 0.4 | 0.3 |
| Bio08 | Mean Temperature of Wettest Quarter | 1.9 | 0.8 | 1.5 | 1.9 | 0.7 | 1.7 |
| Bio09 | Mean Temperature of Driest Quarter | 0.4 | 0.5 | 0.5 | 0.4 | 0.6 | 0.4 |
| Bio15 | Precipitation Seasonality | 7.1 | 7.3 | 7.7 | 6.7 | 8.0 | 7.2 |
| Bio18 | Precipitation of Warmest Quarter | **55.1** | **55.1** | **55.6** | **54.3** | **55.4** | **56.3** |
| Bio19 | Precipitation of Coldest Quarter | **22.2** | **23.3** | **21.5** | **23.8** | **23.4** | **22.5** |
| *AUC* |  | *0.975* | *0.973* | *0.974* | *0.975* | *0.973* | *0.973* |
