## Supplementary Material 3 for "Evolutionary history of *Castanea sativa* Mill. in the Caucasus driven by Middle and Late Pleistocene paleoenvironmental changes"

ORCID:

BB: 0000-0003-4189-5392

KS: 0000-0002-1830-0816

ŁW: 0000-0002-4060-9801

PT: 0000-0003-3115-3301

GK: 0000-0002-5654-4061

VF: 0000-0002-4004-234X

AB: 0000-0002-4629-1507

MD: 0000-0001-7017-5541

**Table S1.** Characterization of nuclear microsatellites used in the study

| Loci | Number of alleles | *Ho* | *uHe* | Fis | Null | Observed range | Motif | Forward and Reverse primer (5′-3′) | Reference |
| --- | --- | --- | --- | --- | --- | --- | --- | --- | --- |
| EMCs15 | 6 | 0.475 | 0.482 | -0.002 | 0.022 | 74-95 | $({CAC)}_{9}$ | F _ CTCTTAGACTCCTTCGCCAATC R_ CAGAATCAAAGAAGAGAAAGGTC | Buck et al. 2003 |
| EMCs2 | 5 | 0.030 | 0.030 | -0.011 | 0.015 | 151-166 | $({CGG)}_{7}$ | F _ GCTGATATGGCAATGCTTTTCCTC R_ GCCCTCCAGCCTCACCTTCATCAG | Buck et al. 2003 |
| CsCAT6 | 14 | 0.708 | 0.721 | 0.001 | 0.020 | 144-176 | $({AC)}_{24}AT{(AC)}_{4}$ | F _ AGTGCTCGTGGTCAGTGAG R_ CAACTCTGCATGATAAC | Marinoni et al. 2003 |
| EMCs13 | 7 | 0.508 | 0.452 | -0.143 | 0.010 | 139-163 | $({GCA)}_{8}$ | F _ TAGTCGGAGTACGGGCACAG R_ TGATATGAGCATTTGACTTTGATT | Buck et al. 2003 |
| CsCAT15 | 10 | 0.555 | 0.560 | -0.009 | 0.017 | 121-143 | $({TC)}_{12}$ | F _ TTCTGCGACCTCGAAACCGA R_ GCTAGGGTTTTCATTTCTAG | Marinoni et al. 2003 |
| CsCAT1 | 19 | 0.310 | 0.404 | 0.218 | 0.089 | 175-229 | $({TG)}_{5}TA{(TG)}_{24}$ | F _ GAGAATGCCCACTTTTGCA R_ GCTCCCTTATGGTCTCG | Marinoni et al. 2003 |
| CsCAT14 | 11 | 0.614 | 0.616 | -0.013 | 0.020 | 133-169 | $({CA)}_{22}$ | F _ CGAGGTTGTTGTTCATCATTAC R_ GATCTCAAGTCAAAAGGTGTC | Marinoni et al. 2003 |
| EMCs22 | 16 | 0.685 | 0.745 | 0.064 | 0.046 | 124-156 | $({GA)}_{19}$ | F _ 6-FAM–GTGCCTCTGTATGCATGGTAAGC R_ CCAGGTTTAAGAAAGCAAGCATAAC | Buck et al. 2003 |
| CsCAT41 | 25 | 0.609 | 0.823 | 0.247 | 0.120 | 206-290 | $({AG)}_{20}$ | F _ AAGTCAGCAACACCATATGC R_ CCCACTGTTCATGAGTTTCT | Marinoni et al. 2003 |
| Mean | 12.56 | 0.50 | 0.54 | 0.04 | 0.04 | ­— | — | — | — |

Marinoni D., Akkak A., Bounous G., Edwards K.J and Botta. R. 2003. Development and characterization of microsatellite markers in *Castanea sativa* (Mill.). Molecular Breeding. 11(2): 127-136.

Buck E.J., Hadonou M., James C.J., Blakesley D and Russell. K. 2003. Isolation and characterization of polymorphic microsatellites in European chestnut (*Castanea sativa* Mill.). Molecular Ecology Notes. 3(2): 239-241.

**Table S2.** The pairwise differentiation (Fst) among 21 studied populations of *C. sativa* in the South Caucasus computed in FreeNA. The significance was assessed based on 9,999 permutations. All values are in the range of 95% CI.

| Fst | LC1 | LC2 | LC3 | LC4 | LR2 | LR1 | WGC1 | WGC2 | WGC3 | WGC4 | WGC5 | WGC6 | WGC7 | CGC1 | CGC2 | CGC3 | CGC4 | EGC1 | EGC2 | EGC3 |
| --- | --- | --- | --- | --- | --- | --- | --- | --- | --- | --- | --- | --- | --- | --- | --- | --- | --- | --- | --- | --- |
| LC1 |  |  |  |  |  |  |  |  |  |  |  |  |  |  |  |  |  |  |  |  |
| LC2 | 0.011 |  |  |  |  |  |  |  |  |  |  |  |  |  |  |  |  |  |  |  |
| LC3 | 0.027 | 0.024 |  |  |  |  |  |  |  |  |  |  |  |  |  |  |  |  |  |  |
| LC4 | 0.024 | 0.036 | 0.004 |  |  |  |  |  |  |  |  |  |  |  |  |  |  |  |  |  |
| LR1 | 0.010 | 0.033 | 0.038 | 0.025 |  |  |  |  |  |  |  |  |  |  |  |  |  |  |  |  |
| LR2 | 0.037 | 0.040 | 0.042 | 0.043 | 0.027 |  |  |  |  |  |  |  |  |  |  |  |  |  |  |  |
| WGC1 | 0.023 | 0.059 | 0.065 | 0.038 | 0.014 | 0.055 |  |  |  |  |  |  |  |  |  |  |  |  |  |  |
| WGC2 | 0.072 | 0.107 | 0.106 | 0.084 | 0.053 | 0.113 | 0.057 |  |  |  |  |  |  |  |  |  |  |  |  |  |
| WGC3 | 0.036 | 0.074 | 0.063 | 0.040 | 0.024 | 0.052 | 0.038 | 0.076 |  |  |  |  |  |  |  |  |  |  |  |  |
| WGC4 | 0.021 | 0.048 | 0.061 | 0.050 | 0.005 | 0.027 | 0.024 | 0.064 | 0.030 |  |  |  |  |  |  |  |  |  |  |  |
| WGC5 | 0.011 | 0.049 | 0.043 | 0.028 | 0.009 | 0.038 | 0.025 | 0.067 | 0.004 | 0.023 |  |  |  |  |  |  |  |  |  |  |
| WGC6 | 0.011 | 0.038 | 0.048 | 0.043 | 0.029 | 0.060 | 0.044 | 0.092 | 0.039 | 0.050 | 0.012 |  |  |  |  |  |  |  |  |  |
| WGC7 | 0.017 | 0.056 | 0.054 | 0.043 | 0.023 | 0.063 | 0.036 | 0.074 | 0.035 | 0.045 | 0.018 | 0.022 |  |  |  |  |  |  |  |  |
| CGC1 | 0.076 | 0.122 | 0.141 | 0.133 | 0.075 | 0.133 | 0.093 | 0.124 | 0.075 | 0.068 | 0.066 | 0.093 | 0.114 |  |  |  |  |  |  |  |
| CGC2 | 0.063 | 0.081 | 0.072 | 0.049 | 0.066 | 0.089 | 0.080 | 0.152 | 0.068 | 0.093 | 0.072 | 0.077 | 0.072 | 0.172 |  |  |  |  |  |  |
| CGC3 | 0.059 | 0.074 | 0.060 | 0.046 | 0.045 | 0.072 | 0.064 | 0.126 | 0.062 | 0.057 | 0.062 | 0.069 | 0.075 | 0.124 | 0.030 |  |  |  |  |  |
| CGC4 | 0.073 | 0.115 | 0.121 | 0.115 | 0.102 | 0.151 | 0.116 | 0.146 | 0.116 | 0.125 | 0.088 | 0.064 | 0.104 | 0.145 | 0.107 | 0.125 |  |  |  |  |
| EGC1 | 0.070 | 0.107 | 0.098 | 0.080 | 0.073 | 0.123 | 0.070 | 0.133 | 0.070 | 0.104 | 0.072 | 0.073 | 0.071 | 0.119 | 0.042 | 0.050 | 0.100 |  |  |  |
| EGC2 | 0.054 | 0.081 | 0.080 | 0.062 | 0.050 | 0.097 | 0.050 | 0.129 | 0.051 | 0.071 | 0.056 | 0.056 | 0.067 | 0.087 | 0.030 | 0.015 | 0.105 | 0.011 |  |  |
| EGC3 | 0.051 | 0.082 | 0.093 | 0.080 | 0.066 | 0.118 | 0.073 | 0.110 | 0.079 | 0.079 | 0.067 | 0.064 | 0.063 | 0.114 | 0.069 | 0.034 | 0.104 | 0.051 | 0.043 |  |
| EGC4 | 0.151 | 0.164 | 0.150 | 0.135 | 0.116 | 0.148 | 0.158 | 0.174 | 0.135 | 0.140 | 0.142 | 0.168 | 0.163 | 0.196 | 0.136 | 0.106 | 0.266 | 0.148 | 0.135 | 0.152 |

**Table S3.** Regional differences in genetic structure parameters computed for the popualtions from the South Caucasus.

| Group of populations | *Mean He* | *Mean Ar* | *Mean Fis* | *Mean Fst* |
| --- | --- | --- | --- | --- |
| LC/LR/WGC/CGC/EGC | **0.610/0.599/ 0.561/0.512/0.505** | **5.65/5.01/4.88/4.14/3.49** | 0.041/0.089/-0.002/0.016/-0.010 | 0.024/0.023/0.053/0.101/0.083 |
| LC/WGC | 0.610/0.561 | 5.65/4.88 | 0.041/-0.002 | 0.024/0.053 |
| (LC, WGC)/CGC/(EGC | **0.579/0.512/0.505** | **5.16/4.14/3.49** | 0.014/0.016/-0.010 | 0.050/ 0.101/0.083 |
| (LC, LR, WGC)/(CGC, EGC) | **0.582/0.508** | **5.14/3.81** | 0.026/0.001 | 0.046/0.095 |

Bolded are significant differences at p<0.001; LC = Lesser Caucasus, WGC – West Greater Caucasus, CGC – Central Greater Caucasus, ECG – East Greater Caucasus

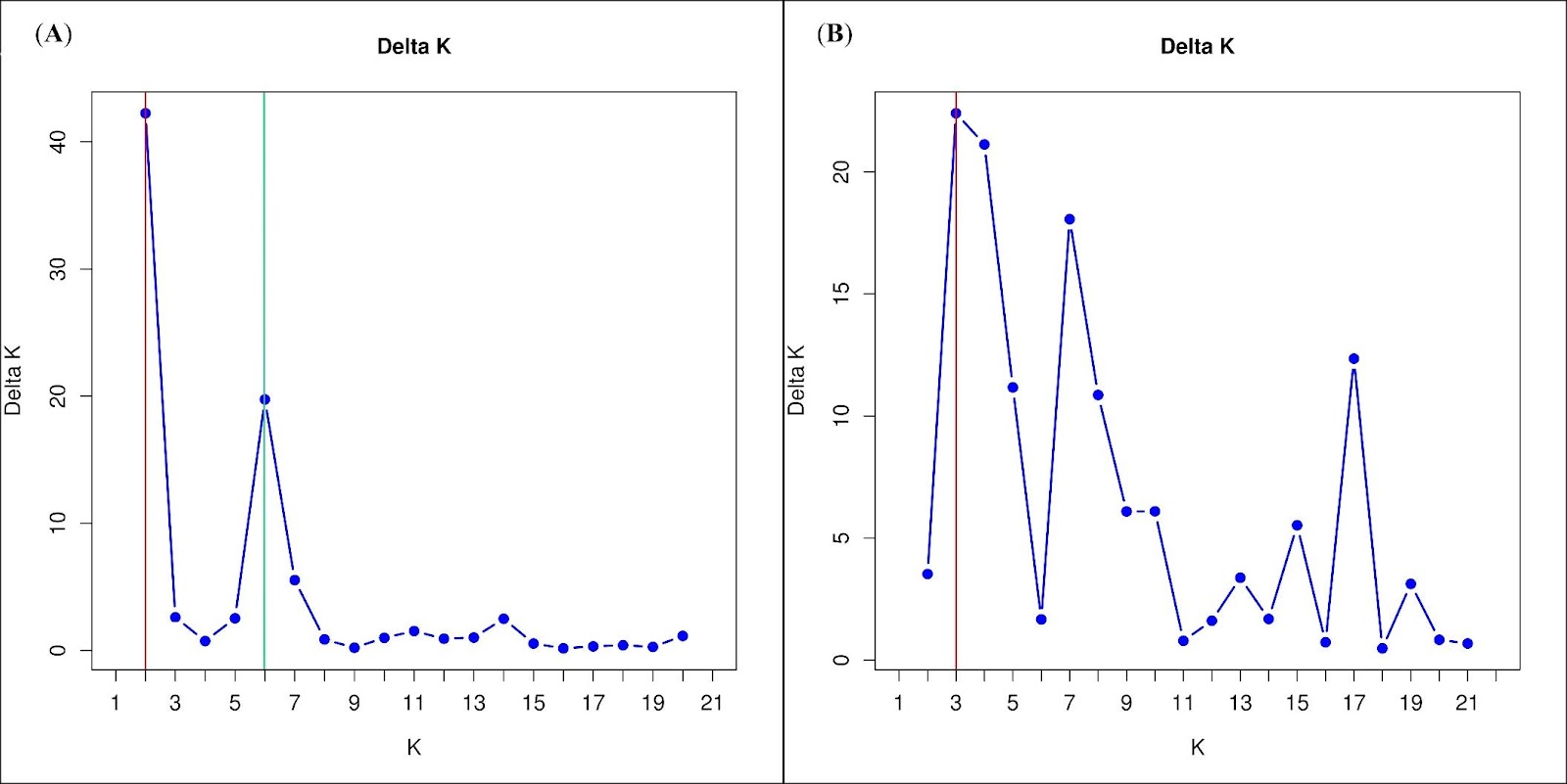

**Fig. S1.** Estimation of the optimal number of genetic clusters of STRUCTURE results based on Evanno’s ΔK method for 22 studied populations of *C. sativa* from the South Caucasus and North Macedonia.

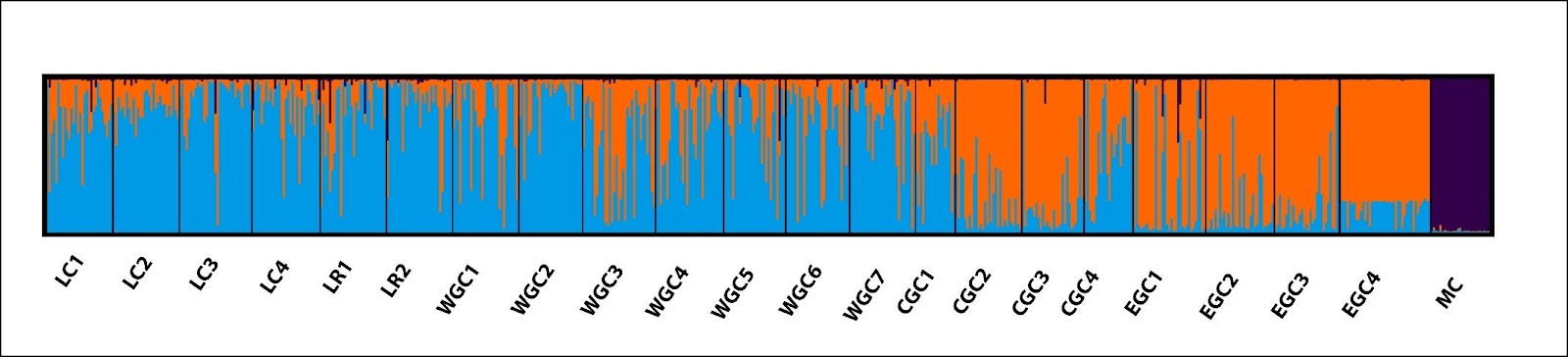

**Fig. S2.** A barplot for K =3 inferred for 22 studied populations of *C. sativa* from the South Caucasus and North Macedonia based on the Evanno’s ΔK method.

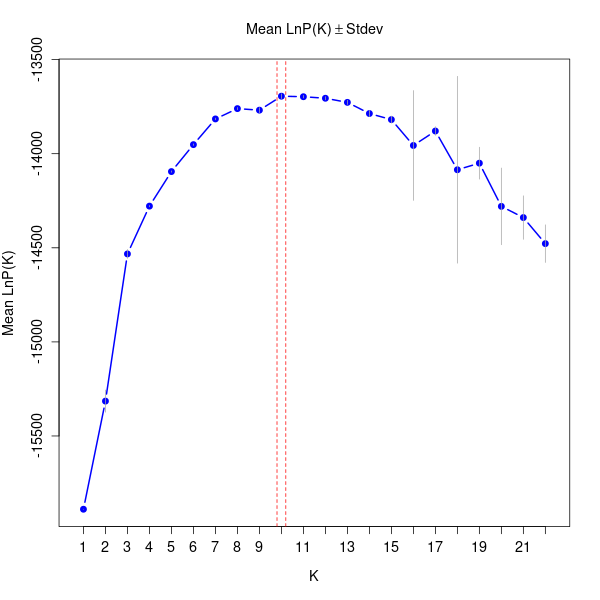

**Fig. S3.**  Estimation of the optimal number of genetic clusters of STRUCTURE results based mean probability method for 22 studied populations of *C. sativa* from the South Caucasus and North Macedonia.

**Fig. S4.** Estimation of the optimal number of genetic clusters of STRUCTURE results based on method of Puechmaille (2016) for 22 studied populations from the South Caucasus and North Macedonia.

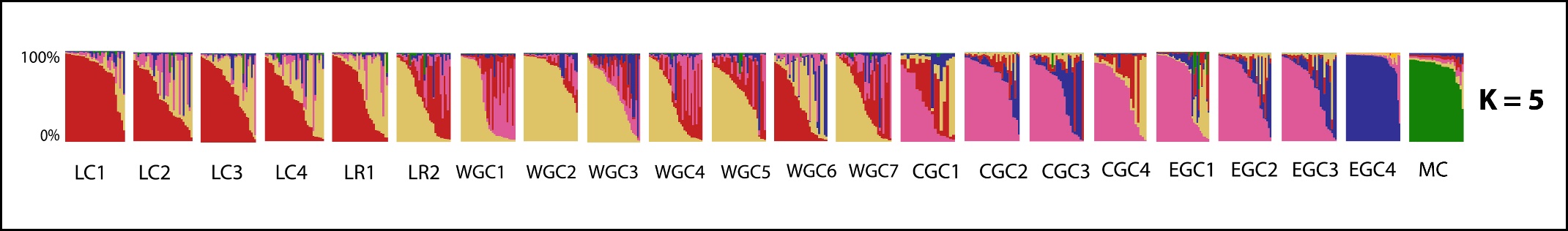

**Fig. S5.** A barplot based on K=5 inferred for 22 studied populations of *C. sativa*  from the South Caucasus and North Macedonia based on an approach of Puechmaille (2016).

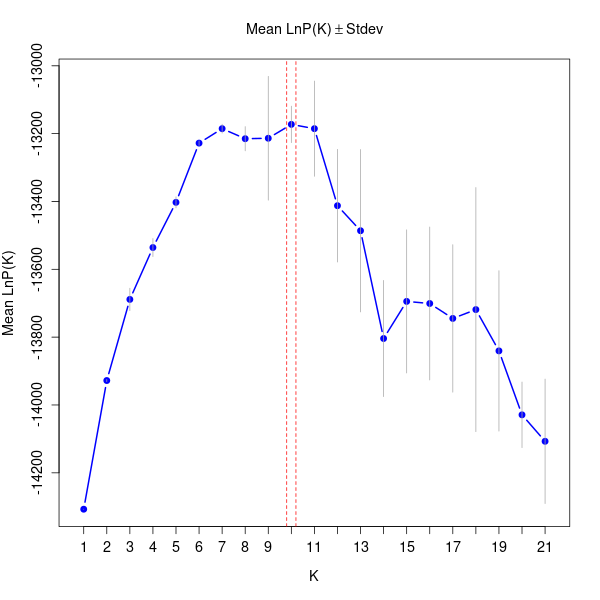

**Fig. S6.** Estimation of the optimal number of genetic clusters of STRUCTURE results based mean probability method for 21 studied populations of *C. sativa* from the South Caucasus.

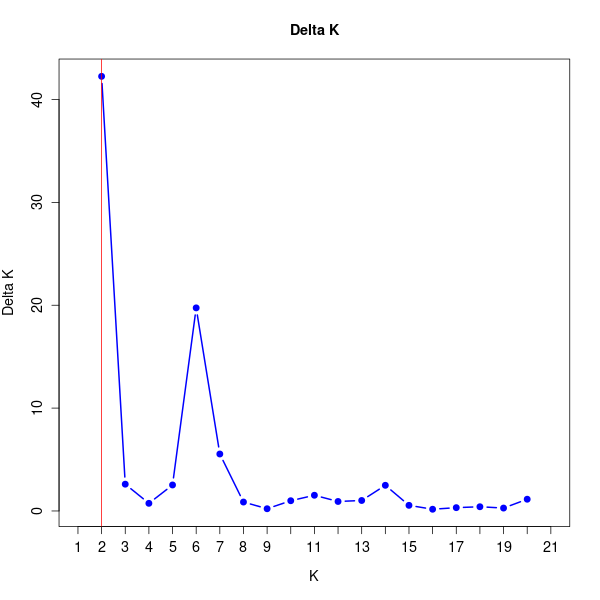

**Fig. S7.** Estimation of the optimal number of genetic clusters of STRUCTURE results based on Evanno’s ΔK method for 21 studied populations of *C. sativa* from the South Caucasus.

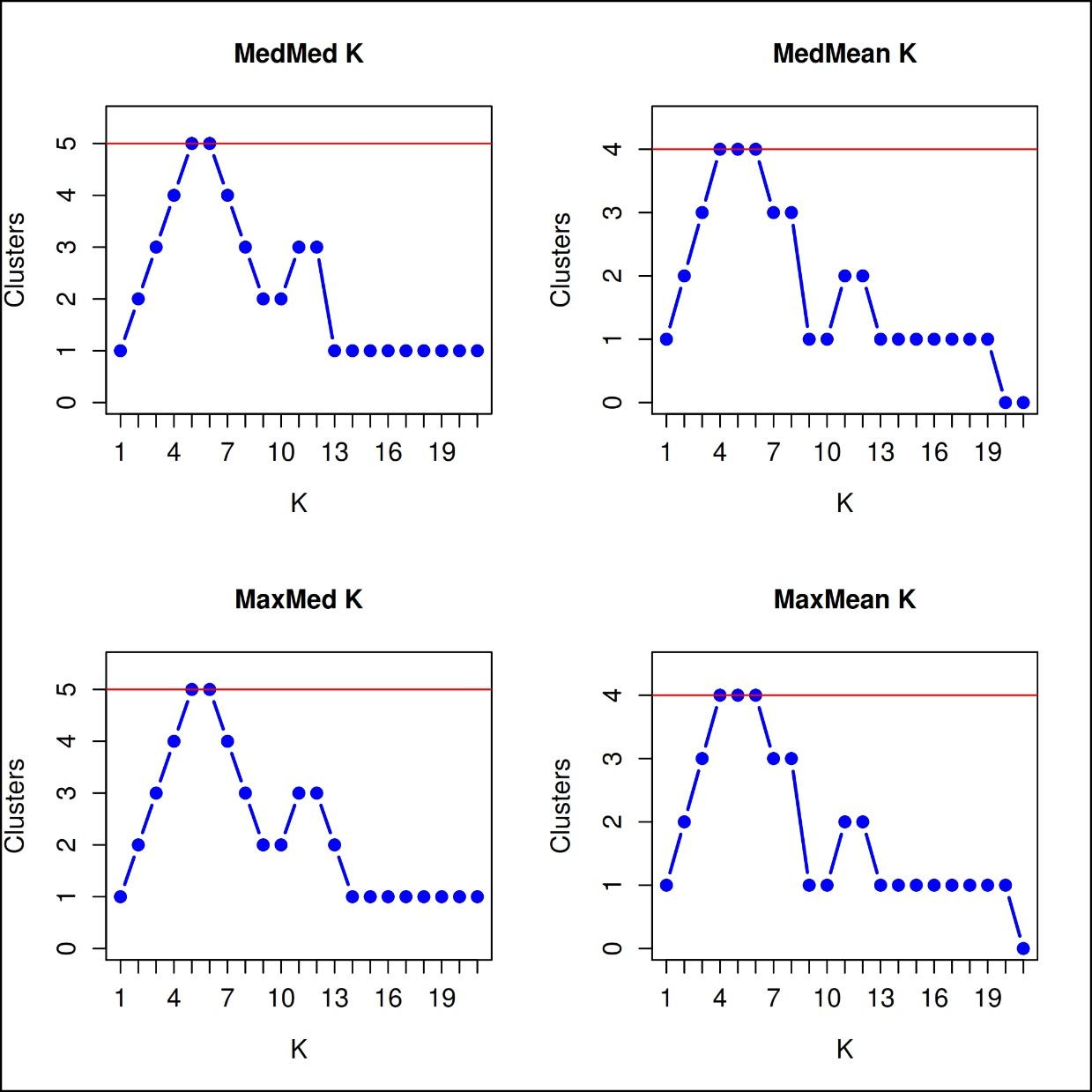

**Fig. S8.** Estimation of the optimal number of genetic clusters for the South Caucasian populations of *C. sativa* (Georgia and Azebijan) based on approach of Puechmaille (2016). The barplot for K=4 was presented in the main text of this article.

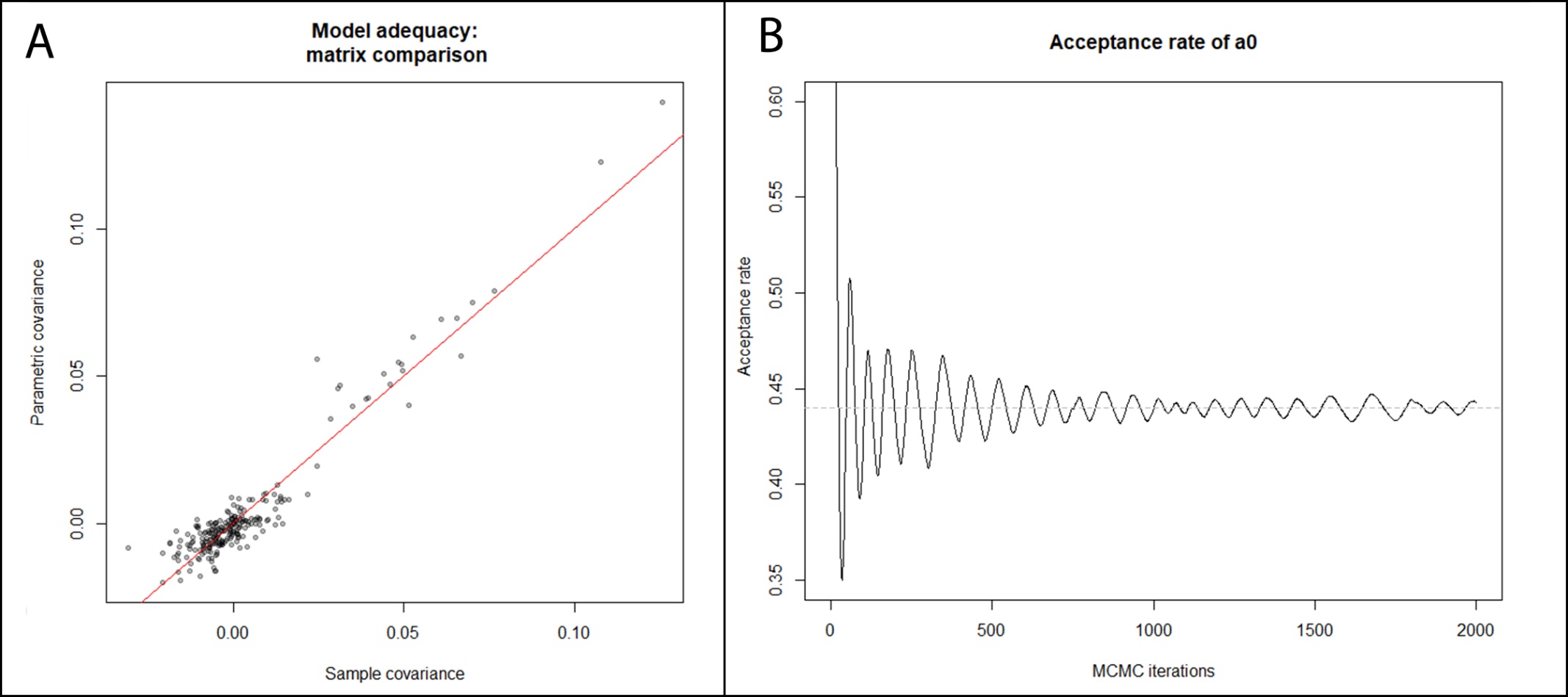

**Fig. S9.** The convergency plots for SpaceMix analysis.
